# Epigenetic reprogramming of melanoma cell state through fatty acid β-oxidation and Toll-like receptor 4 signaling

**DOI:** 10.1101/2022.06.16.496450

**Authors:** Ting-Hsiang Huang, Yilun Ma, Emily Montal, Shruthy Suresh, Mohita M. Tagore, Alexandra Corbin, Dianne Lumaquin, Nathaniel R. Campbell, Arianna Baggiolini, Richard P. Koche, Richard M. White

**Affiliations:** Department of Cancer Biology and Genetics, Memorial Sloan Kettering Cancer Center, New York, NY 10065, USA; Weill Cornell/Rockefeller/Sloan-Kettering Tri-Institutional MD-PhD Program, New York, NY 10065, USA; Cell and Developmental Biology Program, Weill Cornell Graduate School of Medical Sciences, New York, NY 10065, USA; Center for Stem Cell Biology and Developmental Biology Program, Memorial Sloan Kettering Cancer Center, New York, NY 10065, USA; Center for Epigenetics Research, Memorial Sloan Kettering Cancer Center, New York, NY 10065, USA

## Abstract

Tumor cells respond to a host of factors from the local microenvironment. Microenvironmental fatty acids can be used by melanoma cells for fuel, but their effects on transcription and epigenetics are still unclear. Here, we show that the fatty acid β-oxidation (FAO) pathway integrates signaling and epigenetics to drive melanoma progression. Using transgenic zebrafish and human cell lines, we find that octanoate, a medium-chain fatty acid, increases tumorigenesis. Octanoate is metabolized via the FAO/ACLY axis into acetyl-CoA, leading to increased histone acetylation. Transcriptomic and epigenetic analyses demonstrate a convergence of inflammatory gene signatures in octanoate-treated melanoma cells. This signature is mediated by TLR4/MyD88 signaling, which is activated by saturated fatty acids like octanoate. Genetic inactivation of either FAO enzymes or TLR4/MyD88 inhibits alterations in histone acetylation, and rescues octanoate-tumor promoting effects. Together, these data demonstrate clear evidence linking fatty acid metabolism and epigenetics to melanoma pathogenesis through TLR4 signaling.

## Introduction

To cope with nutrient restrictive conditions, cancer cells metabolically adapt to their local tumor microenvironment (TME). For example, melanomas can grow into subcutaneous tissues composed of adipocytes, where they take up fatty acids that promote tumor progression (Clement et al., 2020; Zhang et al., 2018a). In addition to uptake, cancer cells can also increase *de novo* synthesis of fatty acids by upregulating the expression of multiple FA enzymes, transporters, and receptors (Koundouros and Poulogiannis, 2020; Menendez and Lupu, 2007). Elevated serum levels of free fatty acids (FFAs) have been associated with high risk of tumorigenesis (Fan et al., 2021; Liu et al., 2014; Zhang et al., 2020). Given the striking variety of fatty acid chain lengths and saturation, understanding the mechanisms by which they act remains a major challenge. These diverse molecules can function as metabolic fuel, signaling molecules, components of membranes and as substrates for intermediate metabolites (Koundouros and Poulogiannis 2020). Fatty acid β-oxidation (FAO) acts as a central bioenergetic pathway with a series of catalytic enzymes that allows a multi-step conversion of FAs into acetyl-CoA. Acetyl-CoA can then be oxidized in the TCA cycle to produce ATP and citrate. Several lines of evidence report that the FAO rate regulated by carnitine palmitoyltransferase 1 (CPT1) is elevated in many types of cancer and is also highly associated with tumor aggressiveness (Park et al., 2016; Qu et al., 2016; Schlaepfer et al., 2014; Shi et al., 2016; Wang et al., 2018). This indicates that the occurrence of an FA metabolism shift can support tumor progression. In addition to being substrates for metabolism, FAs also act as signaling molecules. For example, saturated FAs have been linked to inflammation through recognition via Toll-like receptors (TLRs), mainly TLR4 and other mechanisms (Fritsche, 2015). This signaling results in cytokine and chemokine release from immune cells that inevitably participate in the construction of the TME. Cancer cells can utilize these signals to stimulate proliferation and create a tumor-supportive environment via exhausting anti-tumor immunity or increasing anti-inflammatory responses (Ilkovitch and Lopez, 2008).

Whereas most studies of FA in cancer focus on long chain species such as palmitate, few have focused on medium-chain FAs (mcFAs) despite these being a common dietary constituent. For example, among these FFAs, higher levels of mcFAs like octanoate have been shown to be associated with significant disease progression in response to chemotherapy (Iemoto et al., 2019). In contrast, mcFAs have also been proposed to have antiproliferative effects *in vitro* (Narayanan et al., 2015). Octanoate has also been shown to affect levels of histone acetylation (McDonnell et al., 2016) and can directly octanylate proteins such as ghrelin (Hosoda et al., 2000; Yang et al., 2008). Given that this fatty acid is a common constituent of foods such as milk and coconut oils (Lemarié et al., 2018), a better understanding of how it affects cancer *in vivo* is needed. Here, using a combination of *in vivo* zebrafish models and *in vitro* human cell lines, we demonstrate that this medium-chain saturated fatty acid (mcSFA) contributes to cancer progression through FAO-induced acetyl-CoA abundance, changes in chromatin architecture, and activation of TLR4 in melanoma. TLR4 signaling provides the specificity of chromatin reprogramming and transcriptional events, which convergently facilitate the release of inflammatory factors to promote tumor progression and metastasis in a fatty acid-laden environment.

### Octanoate, a medium-chain fatty acid, promotes melanoma progression

Fatty acids have been demonstrated to contribute to tumor progression in a variety of settings (Baenke et al., 2013; Currie et al., 2013; Koundouros and Poulogiannis, 2020; Röhrig and Schulze, 2016). Most of these studies have focused on either long chain fatty acids like palmitate (Pascual et al., 2017, 2021), or short-chain fatty acids typically produced by the microbiome (Coutzac et al., 2020; Louis et al., 2014; Matsushita et al., 2021). In contrast, medium-chain fatty acids, which are a common dietary constituent, have received less attention. Some studies have suggested that they can be either tumor promoting (Iemoto et al., 2019; Wang et al., 2020) or tumor protective (Lappano et al., 2017; Narayanan et al., 2015; Otto et al., 2008; Wang et al., 2020). To resolve this, we used a rapid transgenic zebrafish model of melanoma in which we could test the effects of this fatty acid class. We chose octanoic acid, an 8-carbon saturated fatty acid, since it is a major constituent of foods, commonly eaten in Western diets, such as milk and coconuts. To determine the effect of octanoate, we used the previously described Transgene Electroporation of Adult Zebrafish (TEAZ) approach (Callahan et al., 2018) to generate aggressive melanoma *in vivo*. This method produces tumors with the genotype of *mitfa*:BRAF^V600E^; *tp53^-/-^*;*ptena/b^-/-^*in the otherwise transparent *casper* background. We administered octanoate by weekly injections under the TEAZ-induced lesions and traced melanoma progression over time (**Fig.1A**) via quantification of tdTomato and melanin area. Octanoate caused a significant increase in tumor burden compared to controls (8.6 mm^2^ in PBS vs. 16.3 mm^2^ in octanoate at the 12^th^ wpe, p=0.0236, **Fig.1B-C**). To test whether this also occurred in human melanoma cells, we treated a panel of cell lines *in vitro* with octanoate across a range of concentrations. While there was variation from line to line, overall, we found an increase in melanoma proliferation when cells were grown in glucose-free medium (**Figs.1D-G**) or complete medium (**Ext. Fig.1**) with low to moderate doses of octanoate (0.1-1 mM). Interestingly, this growth advantage was largely mitigated when cells were grown in glucose-replete conditions, which may be due to glucolipotoxicity as previously observed (Kim and Yoon, 2011; Poitout et al., 2010). These data indicate that the mcSFA octanoate strongly promotes both zebrafish and human melanoma cell growth.

**Figure 1.**
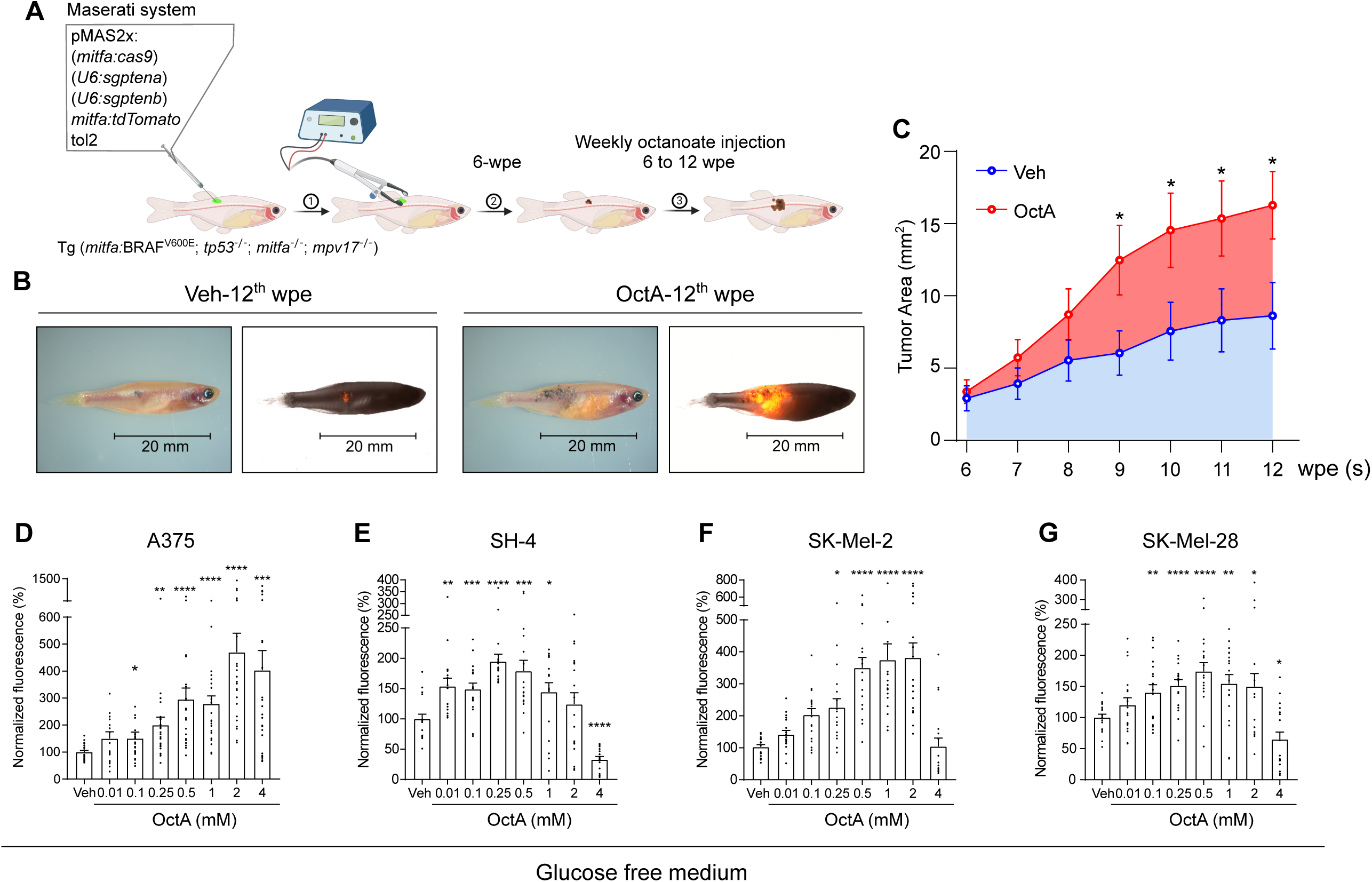
Octanoate stimulates melanoma progression. (**A**) Schematic of TEAZ- and fatty acid injection. Step 1: Plasmids were subcutaneously delivered under fish skin via needle injection followed by electroporation at the injected spot. Step 2: Rescued melanocytes were initially observed in 2-week post-electroporation (wpe) and grew for another 4 weeks to ensure melanocyte expansion. Step 3: After 6-wpe, 1x PBS or 250 µM of octanoate was injected once per week to the fish by syringe needle under melanocytic lesion for 7 weeks. Images were taken from 6 to 12 wpe. (**B**) Color and tdTomato images of representative fish with melanoma at the 12^th^ wpe. (**C**) Quantification of tumor area of tdTomato^+^ and melanin^+^ melanocytes from animals with vehicle or octanoate injection. Veh group was shown in blue (n=33); octanoate (OctA) group was shown in red (n=34). Two biological independent experiments; mean ± SEM, unpaired t-Test, *, p<0.05. (**D-G**) Cell proliferation of human melanomas with the addition of octanoate. 1,500 cells were seeded in each well of 96-well plates and cultured with glucose-deprived DMEM^glucose-(glutaMAX+)^ medium in combination with different concentrations of octanoate for 5 days followed by CyQUANT cell proliferation analysis (Thermo Fisher Scientific #C35011). Mean ± SEM, unpaired t-test, n=4, *, p<0.05; **, p<0.01; ***, p<0.005; ****, p<0.001.

### Octanoate leads to increased global gene expression and chromatin accessibility

To interrogate the mechanism by which octanoate promotes melanoma cell growth, we performed RNA-sequencing of A375 melanoma cells treated with octanoate. We identified 1325 significantly expressed genes (abs (FC) ≥ 1.5, padj ≤ 0.1). GSEA pathway analysis of the genes altered in this dataset were related to defense response to bacterium, lipopolysaccharide (LPS) stimulus, cytokine production, and inflammatory responses (**Fig.2A** and **Ext. Table 1**). In line with these results, the top 10 Gene Ontology Biological Process (GOBP) clusters of identified DEGs are coherently associated with the above bio-themes (**Fig.2B** and **Ext. Table 2**), which are associated with cell-intrinsic inflammatory signal activation.

**Figure 2.**
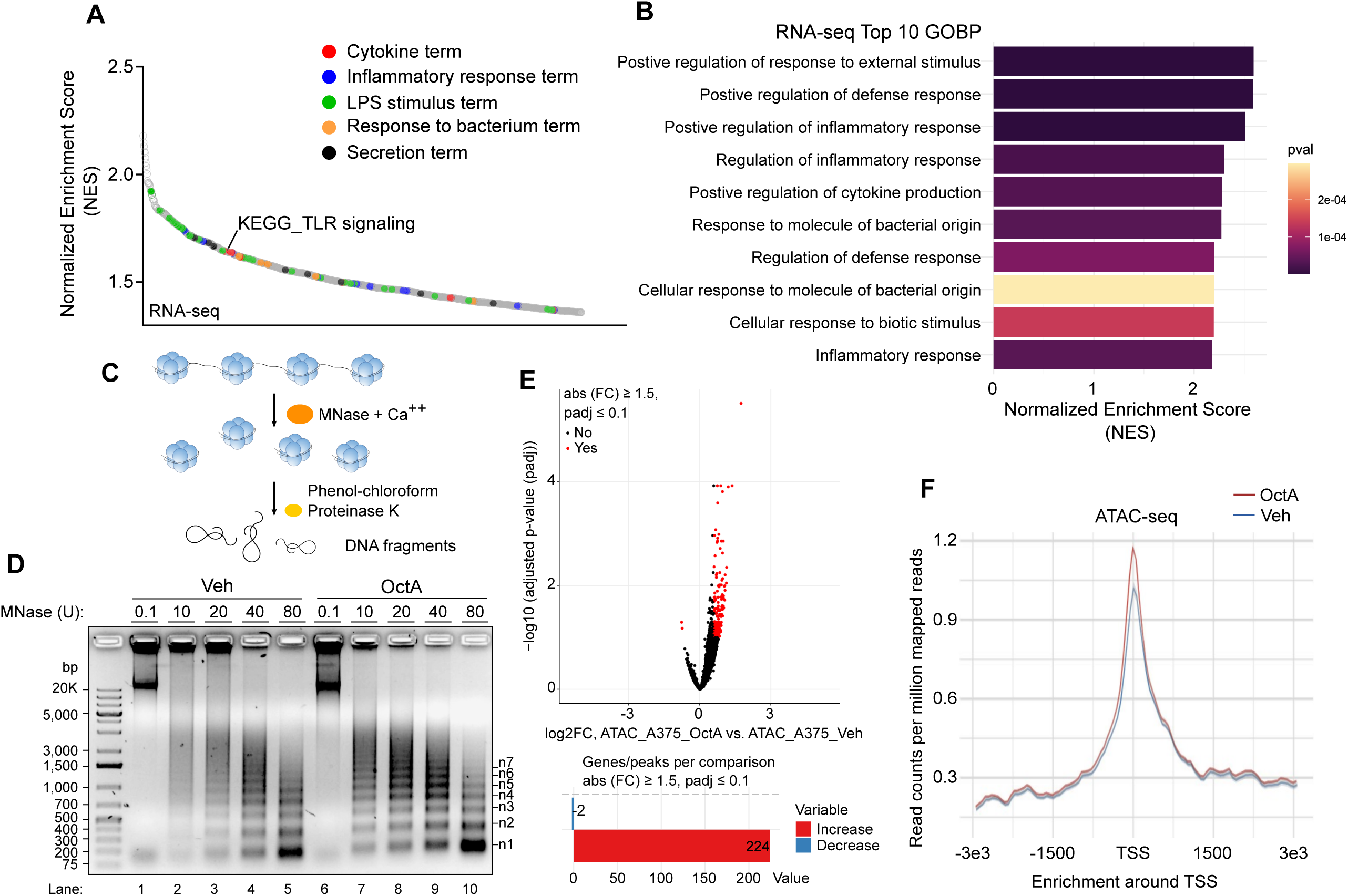
mcFA stimulation leads to enriched inflammatory gene signatures and provides the plasticity of chromatin architecture in melanoma. (**A**) Waterfall plot of all gene sets GSEA analysis in RNA-seq. Top 1,000 pathways categorized to upregulation of cytokine, inflammatory response, LPS stimulus and response to bacterium as well as secretion were highlighted by the color code. (**B**) Top 10 enriched GOBP pathways of RNA-seq responding to octanoate. (**C**) Schematic of Micrococcal Nuclease (MNase) chromatin accessibility assay. (**D**) Size distribution of DNA fragments in 1.2% agarose. n=2. Label on the left, DNA ladder; label on the right, DNA length corresponding to nucleosome number (n). The intensity of DNA fragments was shown in **Ext. Fig. 3**. (**E**) Volcano plot of ATAC-seq analysis. The number of significant peaks were shown in the below bar graph. (**F**) Genetic distribution of mapped ATAC-seq reads around the TSS.

In octanoate treated cells, significantly more genes were upregulated (922 genes) compared to those that were downregulated (403 genes) (**Ext. Fig.2**). Increased gene expression is commonly associated with more open chromatin. To test whether octanoate specifically affected chromatin structure, we used two approaches: the micrococcal nuclease (MNase) assay (**Fig.2C**) and genome-wide ATAC-sequencing. As expected, the MNase treated chromatin fragments showed a size periodicity corresponding to integer multiples of nucleosomes (**Fig.2D**). We found that octanoate-treated cells had a significant increase in open chromatin regions compared to vehicle controls at the same concentration of MNase (**Fig.2D** and **Ext. Fig.3** lane 2 vs. 7, lane 3 vs. 8, lane 4 vs. 9, lane 5 vs. 10). To further test this, we then performed ATAC-seq in the same conditions. This revealed an asymmetric distribution, with 224 significantly increased peaks but only 2 decreased peaks (**Fig.2E**). The regions affected in the ATAC-seq data were associated with less nucleosome occupancy and enrichment of open chromatin at transcriptional start sites (TSS) (**Fig.2F**). This data suggests that mcSFA stimulation leads to decompacted chromatin structure and potentiates transcriptional accessibility.

### Octanoate causes increased histone acetylation

One major mechanism that regulates chromatin compaction is acetylation of histone tails. This was of particular interest to us since it has been previously reported that easily oxidized lipids like octanoate can be metabolized by the fatty acid β-oxidation pathway into acetyl-CoA (McGarry and Foster, 1971), the major cofactor for protein acetylation, including histones (McDonnell et al., 2016). To test this, we added octanoate to a panel of melanoma cells from both zebrafish and humans. This revealed a robust increase in histone abundance at H3 (K9, K27, and pan lysine) and H4 (K12 and pan lysine) in the presence of octanoate compared to vehicle (isopropanol) controls (**Figs.3A-B**; **Ext. Figs.4A-D** and **Ext. Figs.4E-G**). The increase in H3K9ac levels was also visualized by immunofluorescence analysis in zebrafish melanoma Zmel-1 cells (**Fig.3C**). To more directly assess whether lipid-derived metabolites could be utilized in histone acetylation, we measured ^13^C-isotope enrichment of acetylated residues from acidic-purified histones by tracing 1-^13^C-octanoate-derived carbon in A375 cells. Consistent with the previous report (McDonnell et al., 2016), our LC/MS analysis demonstrated that octanoate-derived acetyl groups were incorporated in the core histones including H3K9, K14, K18, and K23 as well as H4K16 at 6-hr or 24-hr octanoate treatment (**Ext. Table 3**). Therefore, this data suggests that fatty acids can be a significant carbon source that allows cells to adapt to nutrient-limited environmental conditions rapidly.

**Figure 3.**
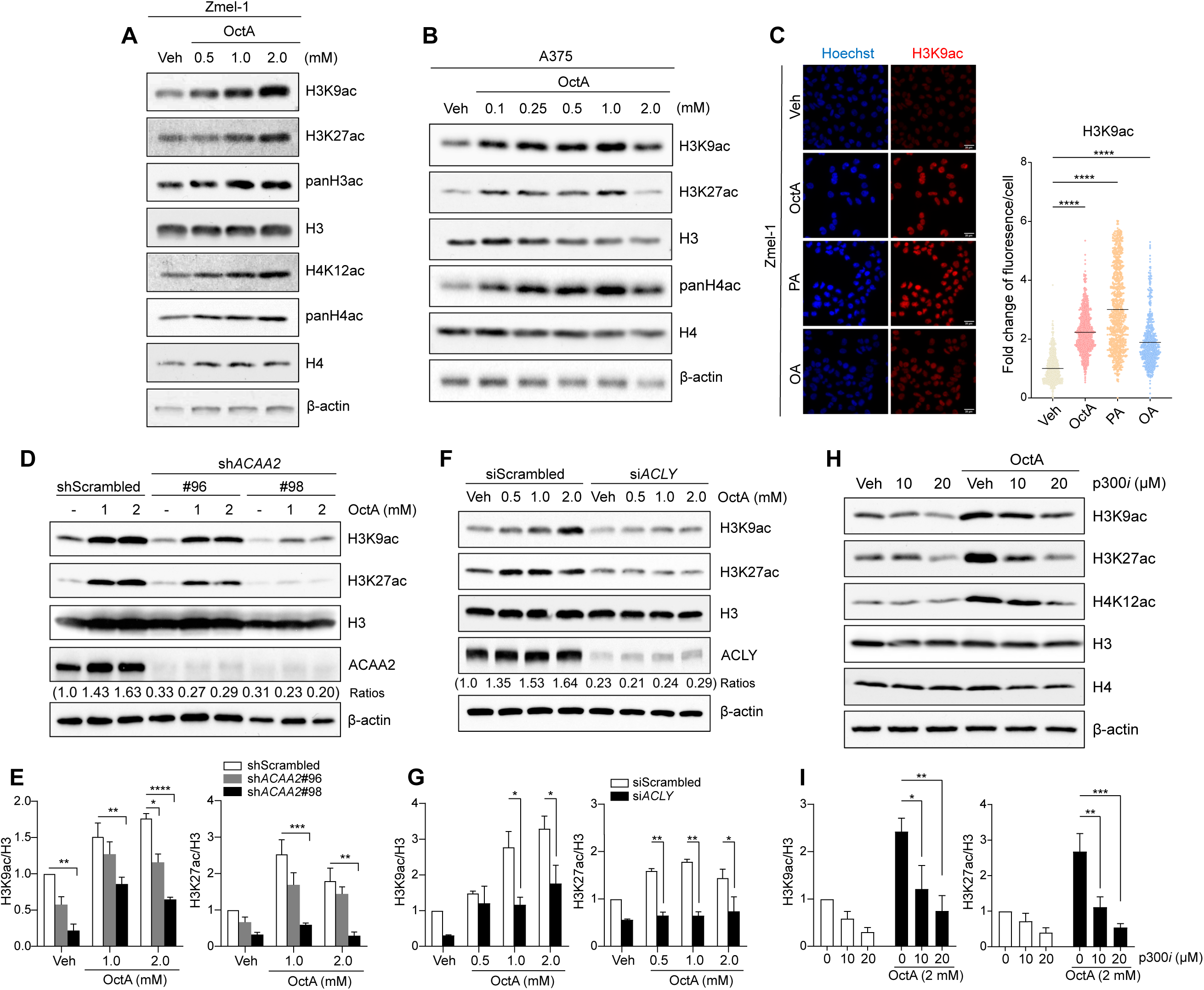
The fatty acid β-oxidation/ACLY/p300 axis regulates the abundance of histone acetylation. Protein levels of histone acetylation in Zmel-1 (**A**) and A375 cells (**B**). Cells were supplied with indicated concentrations of octanoate for 20 hrs followed by Western blot analysis. β-actin was used as the loading control, respectively, n=3. (**C**) Immunofluorescence of H3K9ac in response to various fatty acid treatments. Zmel-1 were incubated with a vehicle control (BSA+isopropanol), 2 mM of octanoate (OctA) or 250 mM of palmitate (PA) or 250 mM of oleate (OA) for 48 hrs upon glucose-deprived DMEM condition (DMEM^glucose-/glutaMAX+^). Images were taken using ZEISS AXIO microscope with AXIOcam 503 mono camera system under 40X objectives across the whole field. H3K9ac fluorescence of each cell was outlined and normalized based on the nuclear area. Over 200 cells of each replicate were measured and quantified using ImageJ. Mean ± SEM, unpaired t-test, n=2, ****, p<0.001. Western blots of H3 acetylation in comparison between *ACAA2* (**D, E**), *ACLY* (**F, G**) knockdowns and scrambled cells. *ACAA2*-ablated A375 cells were cultured in regular complete DMEM with octanoate for 20 hrs. For siRNA-mediated *ACLY* knockdown cells, cells were incubated with glucose-deprived DMEM^glucose-/glutaMAX+^ overnight followed by adding refreshed DMEM^glucose-/glutaMAX+^ with indicated concentrations of octanoate for 20 hrs. Ratios of ACAA2 and ACLY protein expression normalized by β-actin were shown under the blots (n=3). (**E**, **G**) Fold changes of H3K9ac and H3K27ac relative to each vehicle control were determined by ImageJ analysis with 3 independent replicates (H3K27ac/H3 in (**G**), n=2). Mean ± SEM, 2-way ANOVA, *, p<0.05; **, p<0.01; ***, p<0.005; ****, p<0.001. (**H**) Protein levels of H3 and H4 acetylation in the presence of p300 inhibitors (p300*i*). A375 cells were pretreated with C-646, p300 inhibitors for 1 hr prior to the combination with 2 mM of octanoate in regular DMEM for 20 hrs. (**I**) Quantification results of acetylated H3 on K9 and K27 were shown. Mean ± SEM, 2-way ANOVA, n=3, *, p<0.05; **, p<0.01; ***, p<0.005.

### The fatty acid β-oxidation/ACLY/p300 axis regulates increased chromatin accessibility

We next wanted to test whether fatty acid β-oxidation of octanoate to acetyl-CoA is required for this increase in histone acetylation. We knocked down *HADHA*, the α subunit of the mitochondrial trifunctional enzyme responsible for catalyzing the last three steps of reactions in FAO (Djouadi et al., 2016; Rector et al., 2008), and then measured the effect of octanoate. *HADHA* knockdown caused a ∼50% reduction of FA-driven hyperacetylation at K9 and K27 compared to that in scrambled controls (**Ext. Fig.5**). We additionally knocked down another FAO enzyme, *ACAA2*, which participates in the final catabolic step of FAO (Zhang et al., 2018b). Similar to the effect of HADHA, this led to decreased levels of H3K9ac and K27ac (**Figs.3D-E**). Previous studies have shown that mitochondrial acetyl-CoA transport mediated by ACLY is a prerequisite for nuclear histone acetylation (Lee et al., 2014; Wellen et al., 2009). To test this requirement, we knocked down *ACLY* and found that this markedly reduced the induction of H3K9ac and H3K27ac upon octanoate stimulation (**Figs.3F-G**). Finally, because CBP/p300 serves as the main acetyltransferase for glucose-derived histone acetylation (Bugyei-Twum et al., 2014; Chen et al., 2010), we also tested whether this enzyme was responsible for fatty acid induced hyperacetylation. Blockade of p300 abolished the increase in H3ac and H4ac in the presence of octanoate (**Figs.3H-I**). This data demonstrates that octanoate-driven histone hyperacetylation is mediated by fatty acid-oxidation and ACLY-dependent metabolic pathways.

### Secretory and inflammatory gene expression is enriched by octanoate

To further interrogate the correlation between histone hyperacetylation and transcriptomic consequences in response to octanoate, we performed H3K27ac ChIP-seq, a marker of active promoters. Consistent with the asymmetric upregulation of genes and increased chromatin accessibility seen above, we found 1303 significant hyperacetylated peaks and only 24 hypoacetylated peaks (abs (FC) ≥ 1.5, padj ≤ 0.1) after octanoate administration (**Fig.4A**). GSEA pathway analysis showed enriched inflammatory and cytokine gene clusters similar to the transcriptomics analysis (**Fig.4B** and **Ext. Table 4**). Inflammatory proteins such as cytokines are often linked to secretory programs, and consistent with this we found several upregulated secretion-associated pathways in both sequencing datasets (**Fig.2A** and **Fig.4B**). To comprehensively profile proteins with secretory propensity from our dataset, we cross-referenced the upregulated genes from RNA-seq and ChIP-seq to the Human Protein Atlas secretome dataset ([The human blood proteins - secretome](https://www.proteinatlas.org/humanproteome/blood+protein/secretome)) (Uhlén et al., 2019). The secretome dataset was filtered by predicted secretion propensity (1755/2793) (Filtered “secreted” in the column BE of **Ext. Table 13**). The RNA-seq and ChIP-seq showed ∼14% (130/922) and ∼11% (123/1098) of genes overlapped with the secretome database, respectively (**Fig.4C**). Overall, ∼21% of the common genes (56/272) in both datasets (**Ext. Table 5**) were found in the secretome (**Fig.4C** and **Ext. Table 6**). Pathway analysis using PANTHER gene ontology (geneontology.org) (Harris et al., 2004) indicated that 47 out of these 56 secreted proteins were characterized into ten subcategories, including cytokines/chemokines/interleukins (IL-1α, IL-11, IL-37, CXCL3, CCL20, CSF20), peptide hormones (ADM2, PTHLH, STC1), growth factors (INHBA, INHBE, BMP2, BMP6, EREG, GDF15), membrane-bound signal molecules (SEMA3A, SEMA3D, SEMA3F), intracellular signaling molecules (ANGPTL4, ANGPT4, VGF, NOG, WNT10B), metalloproteases/proteases (ADAMTS6, ADAMTS9, ADAMTSL1, ADAMSL4, THSD4 (ADAMTSL6), PCSK1), protease inhibitors (FST, SERPIND1, SERPINE1, CPAMD8, TFPI2), extracellular matrix proteins (COL6A3, NTN4, MUC5AC), transmembrane-signaling receptor (IL-1RL1, IL4R, SFRP1) and cell adhesion molecules (NRCAM, LAMA4) (**Fig.4C** and **Ext. Table 6**). These results indicate that melanoma cells could alter their secretome in response to mcFA stimulation.

**Figure 4.**
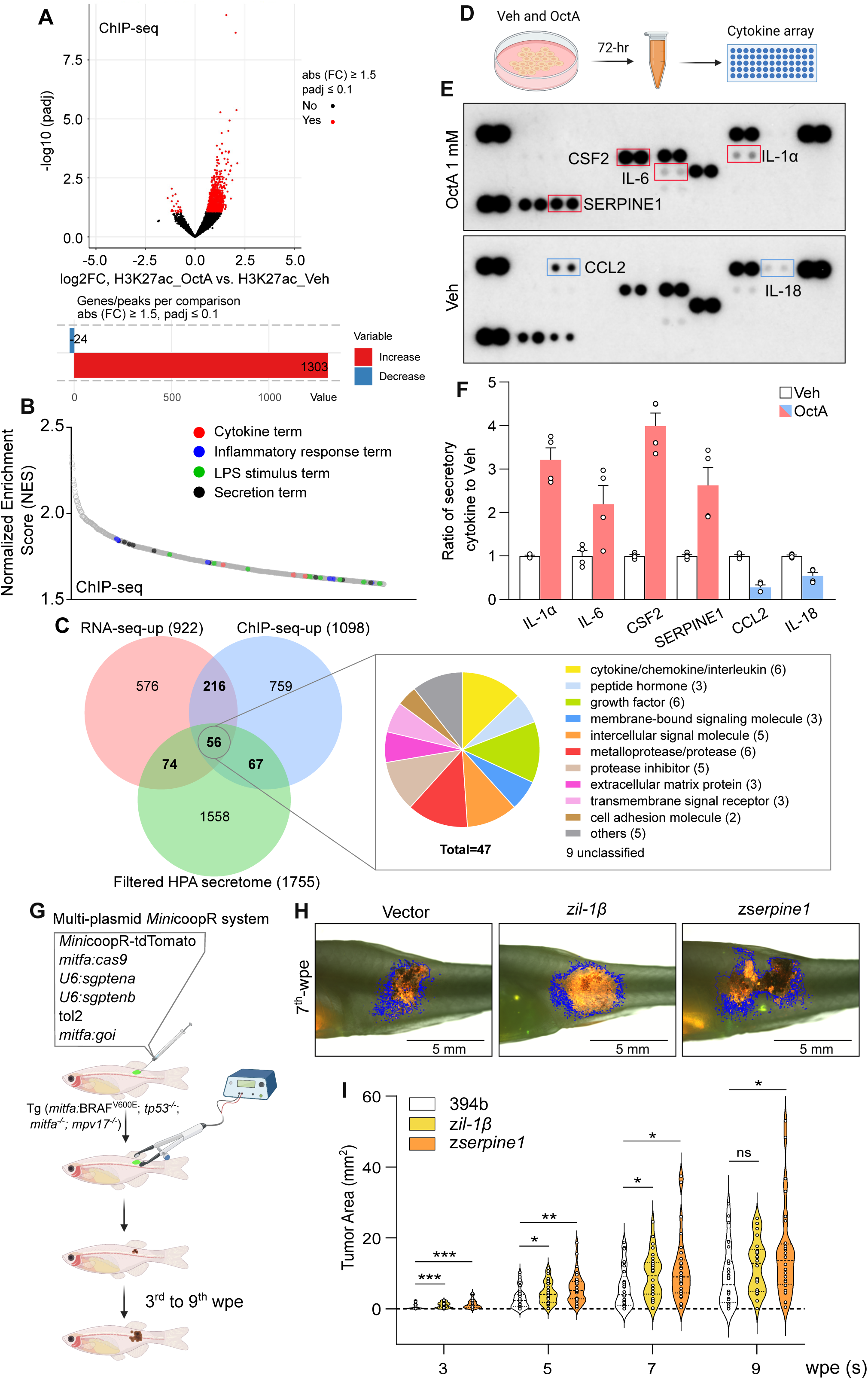
Octanoate enriches secretory and inflammatory gene expression that promotes melanoma progression. (**A**) Volcano plot of H3K27ac ChIP-seq with significant differential gene expression upon octanoate treatment. (**B**) Waterfall plot of all gene sets GSEA analysis in H3K27ac ChIP-seq. Top 1,000 pathways containing upregulation of cytokine or inflammatory response or LPS stimulus or secretion term were presented by the indicated color code. (**C**) Venn diagram between upregulated DEGs of RNA-seq and ChIP-seq as well as the published available HPA secretome database. PANTHER protein class of the common 56 genes was presented by a pie chart. (**D**) Schematic of cytokine expression array. Made by BioRender. (**E**) Western blots of cytokine arrays with a 20-hr treatment of 2 mM octanoate. Elevated or declined protein levels were marked in red and blue open boxes, respectively. (**F**) Quantitative plots of secretory cytokine levels with or without octanoate treatment. Medium collected from A375 culture plates after incubation with octanoate for 72 hrs were applied to the cytokine array. The rest of the steps were performed based on the manufacturer’s instructions. Mean ± SEM, 2 biological independent replicates. (**G**) Schematic of fish melanocyte rescue in combination with mitfa-driven cytokine expression. Made by BioRender. (**H**) Images of representative fish bearing TEAZ transgenic melanoma. In addition to applying the multiple plasmid mixture for melanoma rescue, the quad fish were electroporated in combination with the vector control or plasmid with *mitfa*:z*il-1β* or *mitfa:*z*serpine1*. Images were taken every other week starting from the 3^rd^ to the 9^th^ wpe. The mean of tumor area was shown in (**I**). 394b (vector) (n=30); z*il-1β* (n=30); z*serpine1* (n=30), two biological independent experiments; mean ± SEM, unpaired t-Test, *, p<0.05; **, p<0.01; ***, p<0.005.

To validate the secreted factors obtained from the above analysis, we collected conditioned media from human A375 melanoma cultures upon stimulation with either vehicle or octanoate for 72 hrs, followed by a multiplex antibody cytokine array for 36 human secreted factors (**Fig.4D**). The assay revealed that some cytokines were constitutively expressed under normal conditions (**Fig.4E**), but that octanoate enhances the expression of secreted pro-inflammatory factors such as IL-1α, IL-6, CSF2, and the cytokine-related protein SERPINE1 (**Fig.4E**, **upper panel** and **Fig.4F**). In addition, we analyzed interleukin-1β (IL-1β) expression, a top-ranked DEG in the RNA-seq (**Ext. Table 7**) using ELISA and found IL-1β was secreted abundantly in both LPS which served as a positive control (Fenton et al., 1987) and saturated FA (octanoate and palmitate)-conditioned media (**Ext. Fig.6**). Together, these data suggest melanoma cells transcriptionally reprogram secretory patterns in conjunction with epigenetic regulation of histone acetylation at specific gene sets upon octanoate stimulation.

### Secretory and inflammatory proteins promote melanoma progression *in vivo*

To validate whether the production of octanoate-induced factors by melanoma contributes to tumor progression, we examined the effects of IL-1 and SERPINE1 *in vivo.* IL-1 has been known as a pleiotropic pro-inflammatory cytokine that encompasses two isoforms of IL-1α and IL-1β and is linked to the growth and invasiveness of tumors but also possesses anti-tumorigenic characteristics, owing to the dependency on the stages of cancers and the producing cell (Baker et al., 2019). SERPINE1, a plasminogen activator inhibitor, contributes to a poor prognosis and the progression of various cancers (Pavón et al., 2016; Sakamoto et al., 2021; Seker et al., 2019; Valiente et al., 2014), yet its relevance in skin carcinogenesis is less explored. We overexpressed the fish orthologues *il-1β* and *serpine1* using the TEAZ transgenic model described above (Callahan et al., 2018) (**Fig. 4G**). Both il-1β and serpine1 overexpression (**Ext. Fig. 7**) in fish melanocytes resulted in accelerated melanoma development (**Figs. 4H-I**). Overall, our data demonstrate the functional contributions of melanoma-derived factors (i.e., IL-1 and SERPINE1) in promoting mcFA-driven tumor progression.

### Fatty acid metabolism couples with TLR4 signaling to enhance expression of secretory/inflammatory genes

The above data suggested that octanoate could induce increases in chromatin accessibility and gene expression at inflammatory/secretory loci, but why this is specific to those loci is unclear. We hypothesized that this was related to signaling induced by this fatty acid, which primes those genes for increased expression. Several studies have shown that saturated fatty acids can induce inflammation in various tissues, mainly through toll-like receptor (TLR) signal transduction (Huang et al., 2012; Milanski et al., 2009; Rocha et al., 2016). Intriguingly, GSEA from both RNA-seq and ChIP-seq showed enrichment in response to bacteria and LPS, the major component membrane of gram-negative bacteria, recognized by Toll-like receptor 4 (TLR4) (Poltorak et al., 1998). Furthermore, saturated fatty acids have been demonstrated to promote inflammatory gene signatures likely by indirect interaction with TLR4 (Erridge and Samani, 2009; Lancaster et al., 2018). A recent study reported that TLR signaling plays a role in increasing histone acetylation via metabolic reprogramming and ACLY phosphorylation in macrophages (Lauterbach et al., 2019). Therefore, we hypothesized that TLR signaling in melanoma could be a key mechanism explaining why certain inflammatory/secretory loci were more affected by hyperacetylation than others. We measured mRNA levels of TLR4 downstream cytokines, *IL-1α, IL-1β,* and *CSF2* upon treatment with TLR signaling inhibitors via quantitative real-time PCR (qRT-PCR). Octanoate significantly increased cytokine expression in melanoma, whereas the TLR4 specific inhibitor TAK-242 abrogated the extent of elevated *IL-1α, IL-1β,* and *CSF2* (**Figs. 5A-C**). Surprisingly, inhibition of interleukin-1 receptor-associated kinase 4 (IRAK4), a kinase associated with myeloid differentiation primary response 88 (MyD88), using CA-4948 showed a significant suppressive effect on *CSF2* production but not on *IL-1* (**Figs. 5A-C**), implying a MyD88-independent pathway could be involved in octanoate-mediated *IL-1* expression.

**Figure 5.**
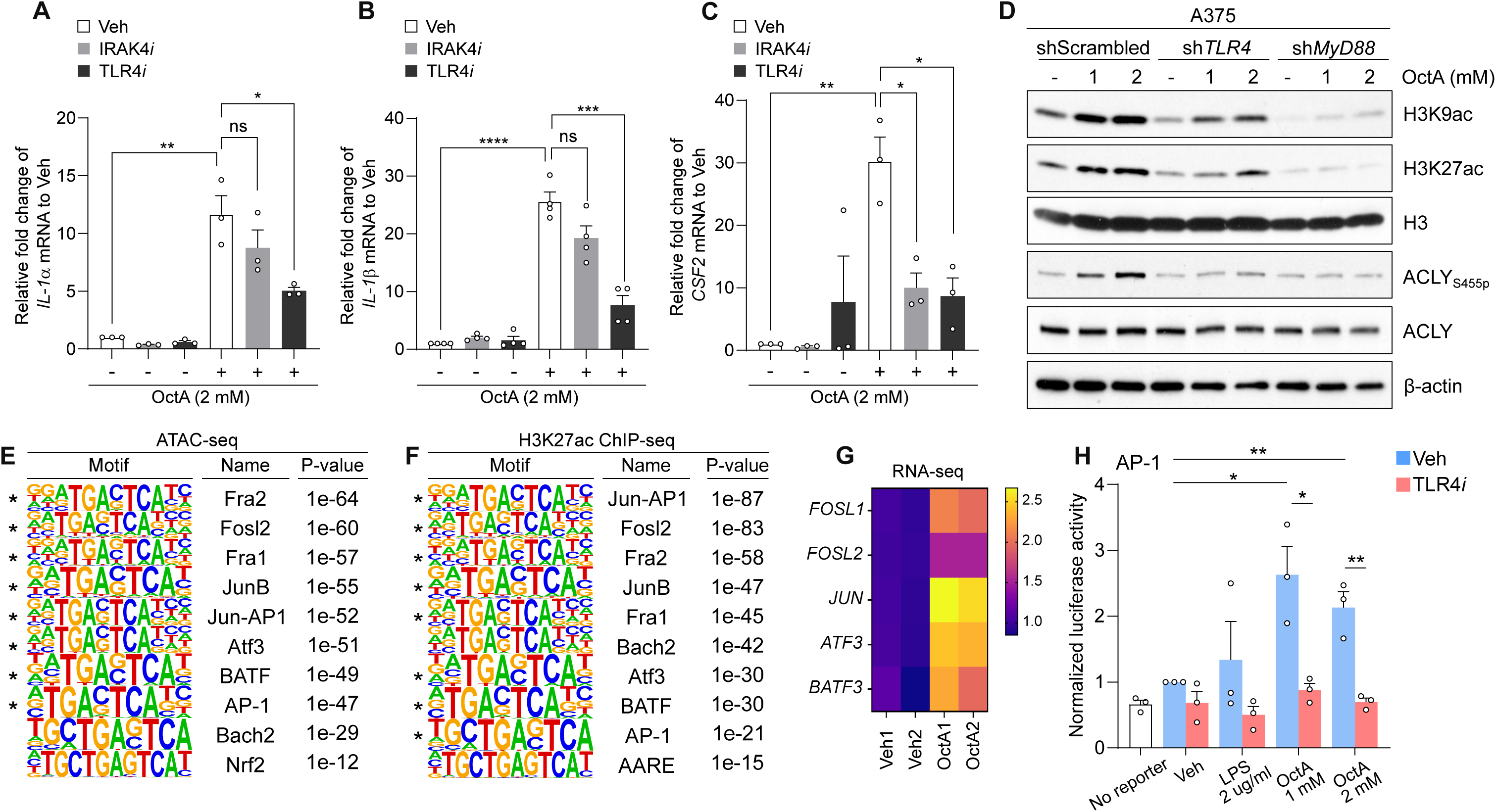
TLR4 signaling offers specificity of fatty acid metabolism for secretory/inflammatory gene expression. Pharmacological inhibition of TLR4 or IRAK4 to examine transcriptional regulation of cytokines induced by octanoate. A375 cells were pretreated with 10 µM of TLR4 inhibitor (TLR4*i*), TAK-242 or 20 µM of IRAK4 inhibitor (IRAK4*i*), CA-4948 for 1 hr prior to a 20-hr co-incubation with octanoate. mRNA levels of *IL-1α* (**A**), *IL-1β* (**B**) and *CSF2* (**C**) were analyzed by qPCR. Mean ± SEM, n=3, unpaired t-Test, *, p<0.05; **, p<0.01; ***, p<0.005; ****, p<0.001. (**D**) Western blot analysis of histone H3 acetylation and ACLY phosphorylation under TLR4 or MyD88 ablation. A375 cells with knockdown of *TLR4* (sh*TLR4*#6897) or *MyD88* (sh*MyD88*#8026) were incubated with octanoate for 20 hrs. Quantification and knockdown results were shown in **Ext. Figs. 8A-C** and **Ext. Figs. 8D-E**, respectively. Top 10 consensus sequences of ATAC-seq (**E**) and H3K27ac ChIP-seq (**F**) by Homer motif analysis. (*AP-1 subunit) (**G**) Heatmap of significant differential expressed AP-1 subunits in RNA-seq (abs(FC) ≥ 1.5, padj ≤ 0.1). The heat color code represents fold changes of normalized reads in each gene between OctA and Veh. (**H**) Luciferase analysis of AP-1 activity in response to LPS or octanoate. Details were described in the Methods. Mean ± SEM, n=3, unpaired t-test, *, p<0.05; **, p<0.01.

Next, we sought to address the effects of TLR4 and its downstream adaptor MyD88 on histone acetylation. Knockdown of *TLR4* using shRNAs inhibited the increase in H3K9ac and H3K27ac compared to the scrambled controls treated with octanoate, while *MyD88*-silenced cells showed an even more potent suppressive effect (**Fig.5D** and **Ext. Figs.8A-B** and **Ext. Figs.8D-E**). Moreover, octanoate-induced ACLY phosphorylation at serine 455 residue (ACLY_S455p_) was attenuated in human A375 melanoma cells lacking *TLR4* or *MyD88* (**Fig.5D** and **Ext. Fig.8C**). These results can be recapitulated by CRISPR-mediated *MyD88* knockout A375 cells (**Ext. Figs.8F-I**) and mouse YUMM3.3 cells upon shRNA-mediated *TLR4* or *MyD88* knockdown (**Ext. Fig.9**). These data are consistent with the idea that the TLR4-MyD88 axis provides specificity for gene expression changes after exposure to octanoate.

TLR4 activates downstream transcription factors such as NF-κB and the activator protein-1 (AP-1) complex, which provides specificity for increased gene expression (Gay et al., 2014; Kawai and Akira, 2007). We reasoned that TLR4, in the context of octanoate, might act through these transcription factors to augment gene expression. We analyzed our ChIP-seq and ATAC-seq data by HOMER and identified dominant consensus sequences of AP-1 components in these datasets, nominating this as a key TF (marked by asterisks, **Figs.5E-F**). We also observed a significant transcriptional upregulation of AP-1 components such as FOSL1, FOSL2, JUN, ATF3, and BATF3 in the octanoate treated cells (**Fig.5G**). To directly test this, we transfected A375 cells with an AP-1 driven luciferase reporter. This showed enhanced AP-1 activity after treatment with octanoate or LPS, and that this activation of AP-1 was suppressed by pharmacological inhibition of TLR4 (**Fig.5H**). Previous work has shown that several kinases such as AKT, JNK, and MEK are upstream mediators of transcriptional machinery in the TLR4 pathway. Inhibition of AKT markedly diminished both octanoate-induced H3K9ac (**Ext. Fig.10A**) and *IL-1α/β* mRNA levels (**Ext. Figs.10B-C**). Similarly, blockade of JNK or MEK also significantly downregulated octanoate-mediated *IL-1α/β* expression (**Ext. Figs.10B-C**). Overall, these data demonstrate that mcFAs activate the AP-1 complex through TLR4 signaling.

### Fatty acid β-oxidation and TLR4 signaling converge to promote melanoma progression *in vivo*

These data indicated that octanoate might converge upon TLR4/MyD88 signaling as well as fatty acid β-oxidation to promote melanoma progression. We compared the invasive capability of mouse melanoma YUMM3.3 cells with knockdown of *Acaa2* or depletion of *Tlr4* and *Myd88* using a 3-dimensional (3D) spheroid *in vitro* invasion assay, which has previously been used to architecturally mimic *in vivo* tumor invasion through the ECM (Vinci et al., 2012, 2015). Control cells initially formed an opaque spheroid in the matrix and spontaneously projected out from the spheroid over time. By day four of octanoate stimulation, the extent of invasion was significantly increased over time compared to vehicle-treated cells (**Fig.6A**). This observation was recapitulated in two other YUMM variants using collagen I-based transwell assays (**Ext. Figs.11A-B**) and is reflected in a migratory gene signature in RNA-seq (**Ext. Fig.11C**). We found that *Acaa2* knockdown decreased both basal as well as octanoate-stimulated invasion in this assay (**Fig.6B**). Even more strikingly, depletion of either Tlr4 or Myd88 eliminated any octanoate induced increase in invasion (**Figs.6C-D**). These *in vitro* results demonstrated that both FA metabolism and TLR4 signaling are critical in octanoate-mediated melanoma invasion.

**Figure 6.**
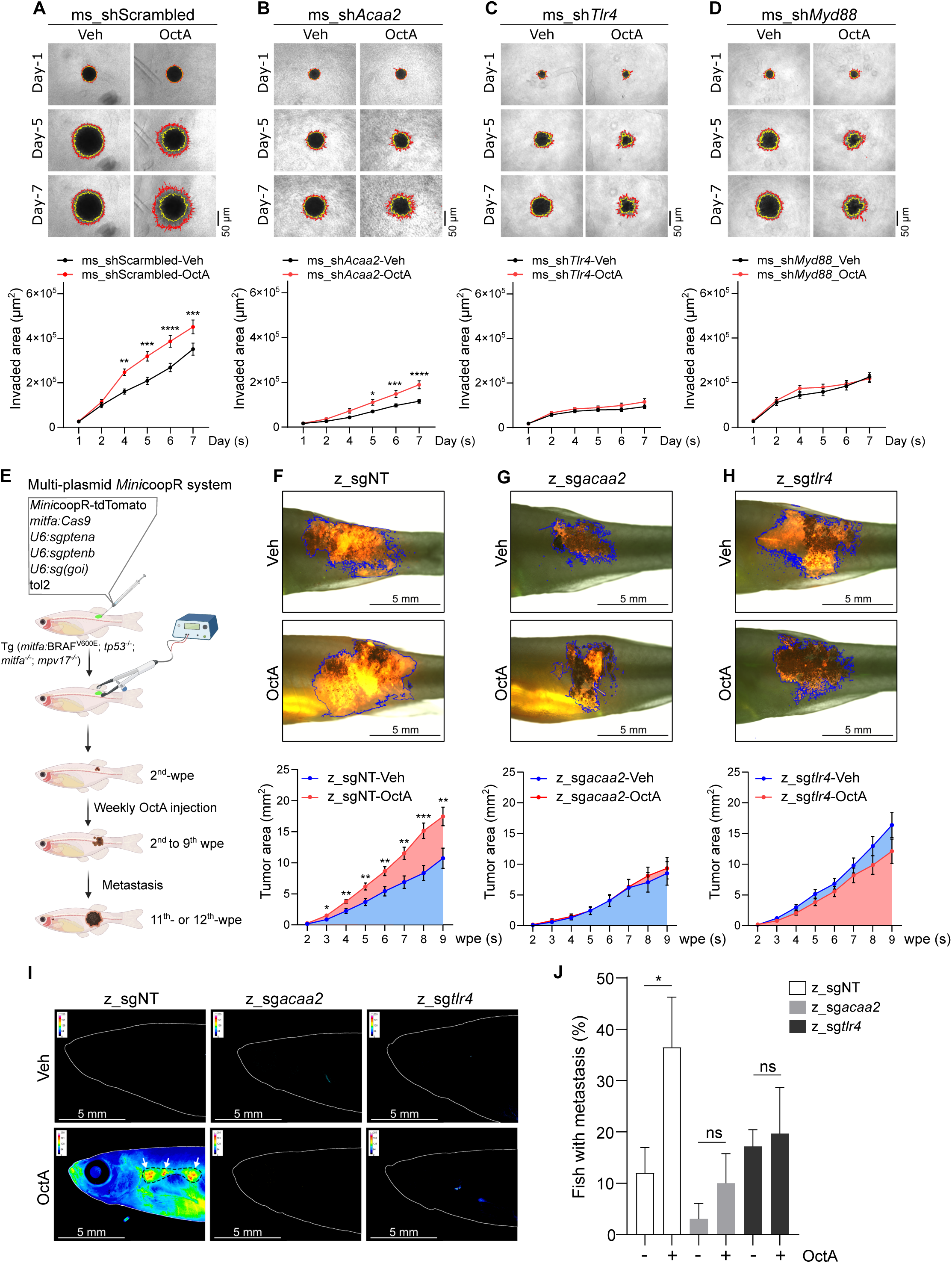
Fatty acid β-oxidation and TLR4 signaling promote fatty acid-mediated melanoma progression *in vitro* and *in vivo*. Comparison of melanoma invasiveness between shScrambled (**A**), sh*Acaa2* (#45) (**B**), sh*Tlr4* (#1488) (**C**) and sh*Myd88* (#7234) (**D**) of YUMM3.3 cells via 3D spheroid *in vitro* invasion assay. Quantification of each knockdown was shown in the lower panel. Mean ± SEM, n=3, unpaired t-test, *, p<0.05; **, p<0.01; ***, p<0.005; ****, p<0.001. (**E**) Schematic of TEAZ transgenic melanoma model in combination with *acaa2^KO^* or *tlr4^KO^* via *Mini*CoopR system. goi: gene of interest. Made by BioRender. Comparison of fish melanoma between Veh and OctA injection upon CRISPR-mediated non-targeting (NT) (**F**) or *acaa2 ^KO^*(**G**) or *tlr4 ^KO^*(**H**) at the 9^th^ wpe. The ROI of the tumor area was highlighted in blue. The quantitative result of each arm was shown in the lower panel. Mean ± SEM, unpaired t-test, *, p<0.05; **, p<0.01; ***, p<0.005. Total fish number, z_sgNT (Veh=53; OctA=57); z_sg*acaa2* (Veh=30; OctA=31); z_sg*tlr4* (Veh=34; OctA=35). n≥3 biological replicates. (**I**) Representative images of metastatic melanoma in knockout fish at 11^th^ wpe. Images were heat-colored by ImageJ to highlight the metastatic area. Zebrafish kidney marrow was marked with a dotted line and melanoma clusters were indicated by white arrows. (**J**) Quantification of fish with metastatic incidence. Mean ± SEM, n≥3, Fisher’s exact test, *, p<0.05, metastasis/total fish number, z_sgNT-Veh (3/34); z_sgNT-OctA (15/43); z_sg*acca2*-Veh (1/26); z_sg*acaa2*-Veh (2/26); z_sg*tlr4*-Veh (5/29); z_sg*tlr4*-OctA (5/27).

Finally, we wanted to test the role of these pathways on tumorigenesis *in vivo*. We utilized TEAZ model to quantitatively track tumor initiation and metastasis. We initiated tumors as described above and used CRISPR to inactivate *acaa2* and *tlr4* (**Fig.6E**). Animals were treated weekly with octanoate injections (**Fig.6F**). Depletion of targets was confirmed by CRISPR-seq (**Ext. Fig.12** and **Ext. Fig.13**). *acaa2^KO^* almost abrogated the effect of octanoate, without affecting vehicle control animals (**Fig.6G**). We found that knockout of *tlr4* slightly increased tumor growth in vehicle treated animals but abrogated the stimulatory effect of octanoate (**Fig.6H** and **Ext. Fig.14**). We continued to monitor these fish until 11-12 wpe when metastases typically appear. Whereas octanoate caused a significant increase in visceral metastases in the sgNT animals, this effect was absent in either *acaa2* or *tlr4* TEAZ-knockouts (**Figs.6I-J**). Taken together, we propose a model (**Ext. Fig.15**) in which fatty acids are metabolized by the fatty acid β-oxidation to promote histone acetylation. This increased histone acetylation amplifies the expression of genes activated by the TLR4/MyD88/AP-1 signaling axis. Together, these act to convergently promote melanoma growth and metastasis.

## Discussion

Metabolic dysregulation is now recognized as a key cancer hallmark that gives tumors the advantage of thriving in nutrient constrained situations. It is increasingly appreciated that cancer cells utilize FA as fuel via intracellular *de novo* FA synthesis or extrinsic FA uptake from the tumor microenvironment to sustain the energy demands for survival (Currie et al., 2013; Koundouros and Poulogiannis, 2020; Röhrig and Schulze, 2016). In addition to being major energy substrates, FAs also function as signaling molecules in various cellular mechanisms taking place in different cell types through G protein-coupled receptors (GPCRs) and toll-like receptors (TLRs). However, most studies are described using long-chain fatty acids (lcFAs) (Hirasawa et al., 2005; Huang et al., 2012; Itoh and Hinuma, 2005; Lee et al., 2003; Milanski et al., 2009; Oh et al., 2010; Shi et al., 2006; Wong et al., 2009). Whether and how other species of FAs play specific roles in cancer progression are minimally reported and thus warrant investigation. To address the question, we utilized an *in vivo* zebrafish transgenic model together with *in vitro* approaches to study the relationship between a dietary mcFA octanoate and melanoma progression. We found that octanoate promotes melanoma cell growth *in vitro* and *in vivo*. When cells respond to octanoate, chromatin becomes more accessible due to abundant acetylated histone decorations mediated through the FAO-ACLY-p300 axis. Transcriptionally, octanoate reshapes melanoma cells, which is in coordination with epigenetic changes of histone acetylation, into an active secretory state that leads to a pro-tumorigenic environment, favoring tumor formation as evidenced by overexpression of IL-1 and SERPINE1 *in vivo*. We observe that TLR4 signaling is required for mcFA-associated acetyl-CoA abundance via ACLY phosphorylation and subsequently induces activation of AP-1 and NF-κB in synergy with the p300 histone acetyltransferase (HAT) machinery at specific genomic loci. Lastly, genetic depletion of TLR4 signaling or impairment of FAO via *ACAA2* knockout suppresses octanoate-induced melanoma development and metastasis.

FA breakdown occurs in mitochondria via FAO to achieve energetic demands and maintain the homeostasis of metabolic networks between various nutrients. Enzymes such as CPT1A and CD36 involved in processing lcFAs for translocation to FAO have been reported in stimulating melanoma progression (Pascual et al., 2017, 2021; Sung et al., 2016). However, it is unclear whether it is also the case for enzymes such as HADHA or ACAA2, which are responsible for the catabolism of FAO. Uniquely, unlike lcFAs that mainly rely on protein-mediated transport and carnitine shuttle system via CPT1 for entering mitochondria, mcFAs can passively diffuse across inner and outer mitochondrial membranes, leading to a rapid FAO. The abundance of acetyl-CoA generated from FAs plays a crucial role in mediating histone acetylation (McDonnell et al., 2016). However, acetyl-CoA’s availability for histone modification depends on the conversion of citrate or acetate to nucleocytosolic acetyl-CoA mediated by ACLY (Campbell and Wellen, 2018; Wellen et al., 2009). Increasing lines of evidence have revealed the multifaceted roles of ACLY in tumorigenesis and DNA repair (Carrer et al., 2019; Lee et al., 2014; Sivanand et al., 2017), which suggests the importance of acetyl-CoA production in cancer cell fates. We show that melanoma cells can employ mcFA-derived acetyl-CoA for global histone acetylation, whereas blockade of the FAO (HADHA and ACAA2)-ACLY axis hinders octanoate-mediated histone acetylation. In addition, octanoate-induced histone acetylation can be blunted by pharmacological inhibition of p300 activity, echoing a recent finding that short-chain fatty acids (scFAs) could directly activate p300 (Thomas and Denu, 2021). However, the selectivity of p300 as a major HAT driven by mcFAs could also be determined by acetyl-CoA concentrations (Pietrocola et al., 2015) or cooperation with its transcriptional cofactors or binding partners. Indeed, we observe octanoate activates AP-1 and NF-κB transcriptional factors that are known for p300 recruitment and interactions at downstream specific genetic loci (Hayden and Ghosh, 2008; Kamei et al., 1996; Vanden Berghe et al., 1999). Other HATs may potentially participate in this process, but they were not investigated due to the significant impacts of p300 on representative H3ac in this study.

Octanoate causes a partially open chromatin status and increases the accessibility of TSS at certain loci. Histone acetylation does not always align with active gene regulation (Xhemalce and Kouzarides, 2010), suggesting that mcFA-associated acetyl-CoA abundance may provide a more specific mechanism to regulate certain genes in melanoma. Similar to this idea, Lee et al. observed that glucose-induced acetyl-CoA abundance causes NFAT1-driven site-specific regulation of histone acetylation and gene expression, which subsequently promotes glioblastoma migration (Lee et al., 2018). Our data provide evidence that profiling octanoate-treated melanoma cells using RNA-seq and ChIP-seq features a high percentage of shared genes in both datasets and convergently demonstrates pro-inflammatory and protein secretion signatures that are regulated by the TLR4 pathway. Besides immune cells, TLR4 is also expressed in various types of cancer cells, and its activation promotes cancer aggressiveness with poor prognosis (Block et al., 2018; Chen et al., 2015; Fukata et al., 2007; Hao et al., 2018; Kim et al., 2012; Wang et al., 2017). For example, a transgenic mouse model demonstrated that the HMGB1 cytokine from UV-irradiated keratinocytes drives TLR4 activation whose downstream inflammatory response plays a critical role in metastatic progression (Bald et al., 2014). Moreover, the level of TLR4 in melanoma is positively correlated with advanced stages of melanoma (Fu et al., 2020; Takazawa et al., 2014). In this respect, we show that octanoate activates AP-1 and NF-κB, and this transcriptional machinery accounts for TLR4-dependent inflammatory gene expression. We also observed that activation of the TLR4-MyD88 pathway contributes to histone hyperacetylation through phosphorylation of ACLY in melanoma, which is consistent with a previous finding in macrophages (Lauterbach et al., 2019). By regulating the availability of nucleocytosolic acetyl-CoA from FAO at the early stage, octanoate-mediated TLR4 signaling directs the HAT transcription machinery potentially composed of the AP-1-NF-κB-p300 complex to the downstream genomic loci for site-specific histone acetylation as shown by enriched H3K27ac.

Consistent with previous evidence showing that long-chain saturated fatty acids (lcSFAs) trigger TLR activation in other cell types (Huang et al., 2012; Milanski et al., 2009; Nicholas et al., 2017; Rocha et al., 2016), we report the application of mcSFA octanoate could mediate TLR4-related downstream response. The mechanism by which TLRs respond to saturated fatty acids (SFAs) is not fully understood. The MD-2-TLR4 receptor complex can recognize LPS by binding lipid A, a lipid endotoxin component responsible for TLR4 stimulation (Park et al., 2009). Given that lipid A variants possess different lengths of acyl chains that determine the immunostimulatory extent of TLR4 downstream events (Maeshima and Fernandez, 2013), it has been widely appreciated due to the structural similarity to lipid A, SFAs could serve as TLR4 agonists or bind with CD14 to regulate CD14-MD-2-TLR4 dimerization docking to the lipid rafts followed by the recruitment of downstream adaptors (Rocha et al., 2016; Wong et al., 2009). However, this concept may depend on the species of SFAs and the number of acyl chains, as a recent discovery demonstrates that palmitate does not have an agonist effect on TLR4 under the experimental settings (Lancaster et al., 2018). Nevertheless, our work provides compelling evidence, for the first time, that mcSFAs can initiate TLR4-mediated inflammatory signals in cancer cells.

The production of inflammatory factors in the TME can either exert their functions on the host immune system or support the proliferation and expansion of cancer cells in both autocrine and paracrine fashion via similar biological processes, which spatiotemporally determines the equilibrium between immunosurveillance and immune escape of melanoma. *In vitro*, we observe that the invasion capability of melanoma can be controlled in both energetic oxidation and TLR4-MyD88 inflammatory signaling fashion. *In vivo*, during melanoma development, we find overexpression of IL-1 or SERPINE1 in melanoma accelerates tumor progression whereas the blockade of FAO or loss of TLR4 ameliorates the extent of mcFA-induced tumor formation and metastatic incidents. We speculate that the prominence of melanoma-secreted immune cytokines in FA-enriched conditions reinforces the development of a tumor-permissive but immunosuppressive microenvironment. This set of diverse secretory factors is intertwined in many aspects, including immunoediting (i.e., IL-1, IL-11, IL-37, CXCL3, CCL20, CSF2), proliferation/ survival of cells (i.e., INHBs, BMPs, EREG, GDF15), angiogenesis (i.e., ANGPT4, ANGPTL4, VGF, ADM2, SERPIND1, SERPINE1), extracellular matrix remodeling (i.e., COL6A3, NTN4, MUC5AC) and tumor metastasis (i.e., NRCAM, LAMA4) via a reciprocal coordinating network to support chronic inflammation and tumor development (Burkholder et al., 2014; Kartikasari et al., 2021). Aside from IL-1 and SERPINE, further investigation using *in vivo* systematic approaches is required to precisely interpret how the other secretory factors we identified are involved in cancer development under the domain of TME.

Of note, TLR4-associated inflammatory response stands at the first host defensive line for the initial construction of innate and adaptive immunity. However, the unresolved inflammation or its activation in cancer cells may cause opposing effects on anti-tumorigenicity depending on the cancer stages and physical conditions such as obesity in the TME (Rogero and Calder, 2018). Several mouse tumorigenic models have shown the essential roles of TLR4 and MyD88 in reducing tumor progression or chemical carcinogenesis via modulating the host immunity (Ahmed et al., 2013; Salcedo et al., 2010; Yusuf et al., 2008), which seem to be contradictory to the growing evidence in the correlation of elevated TLR4/MyD88 with advanced stages and poor prognosis in various cancers (Chen et al., 2015; Fukata et al., 2007; Mittal et al., 2010; Rajput et al., 2013; Rakoff-Nahoum and Medzhitov, 2007). One area for future investigation is the way in which dietary FAs can also directly impact immune cells to enhance immunosuppressive ability through the engagement of FAO in M2 macrophages (Batista-Gonzalez et al., 2019) and myeloid-derived suppressor cells (MDSCs) (Hossain et al., 2015; Veglia et al., 2019). Ultimately, this study demonstrates how FAs can impact melanoma aggressiveness via the integration of epigenetic modifications and inflammatory signaling through TLR4. That improves our current understanding of how the melanoma secretome responds to the tumor microenvironmental influences, leading to advancements in cancer therapy.

## Acknowledgments

We thank the members in the Aquatics Core facility, MSKCC for taking care of animals and their support of this project. We thank Shiva Deljookorani, a former student in the class of 2019 Tri-institutional Gateways Summer Program for helping with this project. At MSKCC, we acknowledge the use of Flow Cytometry Core Facility for FACS experiments, the Integrated Genomics Operations Core for CRISPR-sequencing experiments, and the Gene Editing and Screening Core for IN Cell analysis and shRNA TRC clones. We also thank the Proteomics and Metabolomics Core Facility at Weill Cornell Medicine (WCM) for providing LC-MS services. We thank all the members of White lab for technical support and feedback on the project.

## Funding

This research project was funded by the NIH/NCI Cancer Center Support Grant P30 CA008748, the Melanoma Research Alliance, The Debra and Leon Black Family Foundation, NIH Research Program Grants R01CA229215 and R01CA238317, NIH Director’s New Innovator Award DP2CA186572, The Pershing Square Sohn Foundation, The Mark Foundation for Cancer Research, The American Cancer Society, The Alan and Sandra Gerry Metastasis Research Initiative at the Memorial Sloan Kettering Cancer Center, The Harry J. Lloyd Foundation, Consano and the Starr Cancer Consortium (all to R.M.W.).

T.-H.H. was supported by The Alan and Sandra Gerry Metastasis and Tumor Ecosystems Center Fellowship from MSKCC 2018-2020. Y.M. was supported by the NIH Medical Scientist Training Program Grant T32GM007739 and the Kirschstein-NRSA Predoctoral Fellowship F30CA265124 and the Barbara and Stephen Friedman Pre-doctoral Fellowship from MSKCC 2021-2022. E.D.M. was supported by the NIH F99/K00 Predoctoral to Postdoctoral Fellow Transition Award K00CA223016. S.S. was supported by the Melanoma Research Foundation Career Development Award Grant 719502 and the MSKCC TROT T32 Training Grant T32 CA16000. M.M.T. was supported by Marie-Josée Kravis Women in Science Endeavor Postdoctoral Fellowship and Center for Experimental Immuno-Oncology Postdoctoral Fellowship. A.C. was supported by the SURP program at MSKCC. D.L. was supported by the NIH Kirschstein-NRSA Predoctoral Fellowship F30CA254152 and the NIH Medical Scientist Training Program Grant T32GM007739-42. N.R.C. was supported by the NIH Kirschstein-NRSA Predoctoral Fellowship F30CA220954 and the NIH Medical Scientist Training Program Grant T32GM007739. A.B. was supported by the Swiss National Science Foundation Postdoc.mobility Fellowship P2ZHP3_171967 and P400PB_180672.

## Author contributions

Conceptualization, T.-H.H. and R.M.W.; Methodology, T.-H.H., Y.M. and E.D.M.; Software, R.K., Y.M. and N.R.C.; Investigation, T.-H.H., E.D.M., S.S., M.M.T., A.C. and D.L.; Resources, A.B.; Writing - Original Draft, T.-H.H. and R.M.W.; Supervision, R.M.W.

## Declaration of interests

R.M.W. is a paid consultant to N-of-One Therapeutics, a subsidiary of Qiagen. R.M.W. receives royalty payments for the use of the casper line from Carolina Biologicals.

## Methods

### Resource Availability

#### Lead Contact

Further requests for reagents/materials and information should be directed to Richard M. White.

#### Data Availability

All sequencing data (ATAC/RNA/ChIP-seq) in this study have been deposited and will be publicly available through the NCBI GEO repository. ATAC-seq analysis can be found at GEO (GSE203407). ChIP-seq analysis can be found at GEO (GSE203411). RNA-seq analysis can be found at GEO (GSE203434). CRISPR-seq analysis can be found at GEO (GSE203490). Visualization of H3K27ac ChIP-seq peak distribution (session: H3K27ac_ChIP_seq_octanoate) is publicly available: https://genome.ucsc.edu/s/thuang316/H3K27ac_ChIP_seq_octanoate. Raw counts and differential expression tables of RNA-sequencing and ChIP-sequencing data are shown in the Extended Tables (**Ext. Tables 7** and **14-16**). Raw data of images are available through Mendeley Data (doi: 10.17632/7hbhcn5kbt.1).

### Chemicals

| ***Name*** | ***Manufacture & catalog number*** |
| --- | --- |
| CA-4948 (IRAK4 inhibitor) | Selleck (S9779) |
| C646 (p300 inhibitor) | Cayman Chemical Company (10549) |
| GSK690693 (AKT inhibitor) | Selleck (S1113) |
| Lipopolysaccharides (LPS) | Sigma-Aldrich (L8643) |
| Octanoate | Sigma-Aldrich (C2875) |
| Octanoate-1-13C | Sigma-Aldrich (296457) |
| Oleate | Sigma-Aldrich (03008) |
| Palmitate-BSA | Seahorse Bioscience (102720-100) |
| PD0325901 (MEK inhibitor) | Sigma-Aldrich (PZ0162) |
| SP600125 (JNK inhibitor) | Selleck (S1460) |
| TAK-242 (TLR4 inhibitor) | Sigma-Aldrich (614316) |

### Zebrafish husbandry

Zebrafish stocks were maintained at a density of 5-10/liter under 28°C, light-controlled (14-hr on, 10-hr off) conditions and fed brine shrimp and pelleted zebrafish food. Anesthesia was conducted by Tricaine-S (Western Chemical Inc. #MS-222) with appropriate dilutions from a 4g/L, pH 7.0 stock solution. All fish experiments were performed under the regulation of institutional animal protocols. All the procedures described in the paper were approved by the Memorial Sloan Kettering Cancer Center (MSKCC) Institutional Animal Care and Use Committee (IACUC) with the protocol #12-05-008.

### Zebrafish lines

The transgenic line used in this study was the quad line (*casper* (White et al., 2008) crossing with the triple line (*mitfa*:BRAF^V600E^; *tp53*^-/-^; *mitfa*^-/-^) (Ceol et al., 2011)). Quad fish between 4 to 8 months post fertilization were electroporated to generate melanoma lesions.

### Transgene Electroporation in Adult Zebrafish (TEAZ)

This experimental process was slightly modified based on the previous study (Callahan et al., 2018). To generate BRAF^V600E^; *p53^-/-^*; *ptena/b^KO^* melanomas using Maserati system, the following three tumor initiating plasmids including pMAS2x (Addgene #118844, a gift from Leonard Zon) containing cassettes of *mitfa:cas9*, *U6:sgptena* and *U6:sgptenb* (500 ng/µl), *mitfa:tdTomato* (112 ng/µl) and tol2 (70 ng/µl) were mixed and made up to ∼700 ng/1.5 µl. To generate BRAF^V600E^; *p53^-/-^*; *ptena/b^KO^*melanomas using *Min*icoopR system, multiple plasmids including *Mini*coopR-tdTomato (previously developed in the lab) (350 ng/µl), *mitfa:cas9* (350 ng/µl), *U6:sgptena* (42.5 ng/µl), *U6:sgptenb* (42.5 ng/µl), tol2 (90 ng/µl) were mixed with additional plasmids containing gene of interest (goi) for overexpression (*mitfa:(goi)*) or knockout (*U6:sg(goi)*) and resuspended in sterile water for a final ∼1000 ng/1.5 µl DNA amount prior to electroporation (**Ext. Table 8**). For knockout experiments, two single-guide (sg) plasmids were used to target the same gene. 1.5 µl of plasmid mix was injected using a pulled glass micropipette in the skin between the dorsal fin and the middle line of anesthetized quad fish. Injected fish were put ventrally and immediately electroporated via a platinum probe (A positive electrode was placed on the side of the injected spot.) at 40 volts, 5 pulses. Melanocyte rescue usually occurs in a few weeks post-electroporation on successfully electroporated fish.

### *In vivo* Octanoate administration

When rescued melanocytes were visible to the naked eye (at 6-wpe for the Maserati system, 2-wpe for the *Mini*CoopR system), 250 µM of octanoate or 1x PBS (vehicle) were injected under the electroporated patch once per week in anesthetized TEAZ fish using a Hamilton syringe tipped with a fixed 26s-gauge, bevel needle (Sigma #20734). Fish were then imaged by a Zeiss AxioZoom system with tdTomato and brightfield as well as GFP and mCherry channels (background controls) after injection. The tumor area was measured by identifying the ROIs (region of interest) of all tdTomato sections using the triangle thresholding algorithm implemented in ImageJ v1.53p and then by looking for pigmented tumors in the compatible brightfield images. The tumor area was defined as the mean of total tdTomato^+^ segments including pigmented areas per fish. To determine metastatic incidences, remaining fish were imaged distantly on their anterior parts from the snout to the kidney marrow, where they displayed disseminated tdTomato^+^ melanomas at the 11^th^ or 12^th^ wpe. In addition, GFP signals were subtracted from tdTomato segments, after which all images were uniformly thresholded. The total area of tdTomato^+^ fluorescence ≥ 0.1mm was considered a likelihood for metastasis in animal samples.

### Cell culture

All human melanoma cell lines A375, SH-4, SK-Mel-2, SK-Mel-28 were maintained in DMEM (Invitrogen #11965-118) supplemented with 10% fetal bovine serum (FBS) (Seradigm) at a 37°C incubator with 5% CO_2_. Mouse melanoma YUMM3.3 and 4.1 cells (Meeth et al., 2016) were grown in Gibco DMEM/F-12 (Thermo Fisher Scientific #11320-033) supplemented with 10% FBS at a 37°C incubator with 5% CO_2_. Zebrafish melanoma cell line Zmel-1 constitutively expressing *mitfa*-driven GFP (Heilmann et al., 2015) was grown in DMEM with 10% FBS and glutaMAX (Gibco #35050-061) at a 28°C incubator containing 5% CO_2_.

### Antibodies

All antibodies used in this study were shown in **Ext. Table 9**.

### Short hairpin RNAs (shRNAs)

All shRNAs used in this study were shown in **Ext. Table 10**.

### Primers

All primers and sgRNAs used in this study were listed in **Ext. Tables 11 and 12**.

### Western blot analysis

6.0 x 10^5^ cells were seeded in 6-well plates upon various experimental conditions. Collected cells from plates using cell scrapers (Thermo Fisher Scientific #08-771-1A) were resuspended in 100 ml of 1x prechilled RIPA lysis buffer (Thermo Fisher Scientific #89901) with 1x Halt protease and phosphatase inhibitors (Thermo Fisher Scientific #78443) followed by homogenizing using Branson digital 45 cell sonifier at 10% output, 1-sec burst and 1-sec pause for 15 cycles. After a 10-min centrifuge at 14,000 rpm, 4°C, protein supernatants were collected, and 2 µl of the lysates were subject to 1 ml of 0.2x Bradford protein dye reagent (Bio-Rad #5000006) for protein quantification. 20 µg of protein lysates boiled with SDS sample buffer (Fisher Scientific #NC9140746) were loaded to 4-15% Mini-PROTEAN TGX Precast Protein Gels (Bio-Rad #4561085) for electrophoresis at 100-120 volts followed by semi-dry electrophoretic transfer to nitrocellulose membranes (Bio-Rad #1704270) via Bio-Rad Trans-Blot Turbo transfer system. After 1-hr blocking in 5% nonfat milk/TBS-T (0.1% Tween-20) at room temperature, membranes were incubated with primary antibodies on a shaker, at 4°C, overnight. Membranes were kept in 5% nonfat milk/TBS-T with diluted secondary antibodies on a shaker at room temperature. Following secondary antibody incubation, all blots were washed 3 times with TBS-T (5 mins for each wash) before and after the process. The ECL substrate (Promega #W1001) was dropped widely on all membranes and left for 1 minute, followed by film exposure (Santa Cruz #sc-201697) of the blots. The protein expression level was quantified using ImageJ v1.53p software.

### Immunofluorescence analysis

Melanoma cells were seeded in flat-bottomed 96-well plates overnight. After treatment, cells were fixed with 4% paraformaldehyde for 10 mins and permeabilized in 1x PBS with 0.1% Triton X-100 for another 10 mins. Following a 30-min blocking step with 2% BSA (bovine serum albumin) (Sigma-Aldrich #B4287) resolved in 1x PBS, cells were incubated with 100 µl/well of diluted H3K9ac on a shaker at 4°C overnight and then treated with Alexa Fluor 488 conjugate secondary antibody (Cell Signaling #4412) at room temperature. The images were taken from a mono camera system equipped with an AXIOcam 503 under 40X objectives across the field and with corresponding channels. To quantify histone acetylation levels per cell, ROIs of DAPI-stained nuclei were applied to the images of H3K9ac. The measurement of fluorophore intensity in the ROI was then performed using ImageJ.

### Assay for Transposase-Accessible Chromatin (ATAC)

4.0 x 10^5^ human A375 cells in 6-well plates were treated with vehicle (isopropanol) or 1 mM of octanoate for 48 hrs. After treatment, 1.0 x 10^5^ cells were collected per condition and ATAC-seq was performed as previously described (Buenrostro et al., 2013) with some modifications. Briefly, freshly harvested cells from each condition were lysed in 50 µl ATAC lysis buffer (Tris 10 mM pH 7.4, 10 mM NaCl, 3 mM MgCl_2_, 0.1% NP-40) and incubated with Illumina Tn5 transposase at 37⁰C for 30 mins. Samples were purified using Agencourt AMPure XP beads following which barcoding and library generation were performed using the NEBNext Q5 Hot Start HiFi PCR Master Mix with PCR amplification for 12 cycles. The size selected libraries were run on a HiSeq2500 to get an average of 40-50 million paired end reads per sample.

### Chromatin immunoprecipitation (ChIP) followed by sequencing analysis

5.0 x 10^6^ cells of human melanoma A375 were seeded into 100 mm dishes and were incubated overnight followed by a 48-hr vehicle or 1 mM of octanoate treatment. Fix cells in 10 ml of 4% paraformaldehyde-PBS solution at room temperature with shaking for 8 mins and then neutralize the solution for 2 mins by adding glycine to a final concentration of 125 mM. Wash cell pellets in 1x PBS followed by resuspension in 1 ml pre-chilled ChIP lysis buffer (50 mM, HEPES-KOH pH7.5, 140 mM NaCl, 1% Triton X-100, 0.1% sodium deoxycholate). After a 5-min incubation on ice, break down the cell membrane using a KIMBLE Dounce 2 ml tissue grinder with 30 strokes followed by a 10-min centrifuge at 1,500 rpm, 4°C. Wash nuclei pellets in 500 µl ChIP lysis buffer. Resuspend pellets with 500 µl of nuclei buffer (2 mM EDTA, 100 mM NaCl, Tris-HCl pH 8.0, 0.5% Triton X-100). Genomic DNA was sheared through Branson digital 450 cell sonifier upon the following condition: 10% output at 10-sec burst/1-sec pause for 30 times; 1-min rest on ice per 5 times. Collect supernatant after a 15-min centrifuge at 13,000 rpm, 4°C. Samples were precipitated with H3K27ac antibody (Abcam #4729) overnight on an end-to-end rotator at 4°C followed by a 2.5-hr incubation with protein G Dynabeads (Thermo Fisher Scientific #10003D). Samples were then sequentially washed with TSE (0.1% SDS, 1% Triton-X 100, 2 mM EDTA, 20 mM Tris pH 8.0) x1 (5-min/wash), TSE250 (with 250 mM NaCl) x2, TSE500 (with 500 mM NaCl) x2, and rinsed with TE buffer x2. DNA fragments were resuspended into 100 µl of elution buffer and de-crosslinked by adding RNase at 37°C for 30 mins and proteinase K at 65°C overnight. DNA purification was performed using phenol-chloroform. The DNA library was prepared using NEBNext Utra II DNA Library Prep with Sample Purification Beads (NEB #E7103S) according to the manufacturer’s instruction followed by Sequencing on the Illumina HiSeq conducted by GENEWIZ, LLC. (South Plainfield, NJ, USA).

### Bulk RNA sequencing sample preparation

5.0 x 10^6^ of A375 cells in 100 mm dishes were incubated with vehicle or 1 mM of octanoate for 48 hrs. Cells were detached using cell scrapers and washed off residual medium with 1x PBS after centrifugation, followed by RNA isolation according to the manufacturer’s instruction (Zymo Research #R1055). Snap freeze lysates on dry ice and keep samples in -80°C. RNA library preparation using the NEBNext Ultra RNA Prep Kit for Illumina and next generation sequencing through Illumina HiSeq were conducted by GENEWIZ, LLC. (South Plainfield, NJ, USA).

### Epigenome data analysis

ATAC and ChIP sequencing reads were trimmed and filtered for quality and adapter content using TrimGalore v0.4.5 (https://www.bioinformatics.babraham.ac.uk/projects/trim_galore), with a quality setting of 15, and running version 1.15 of cutadapt and version 0.11.5 of FastQC. Reads were aligned to human assembly hg38 with version 2.3.4.1 of bowtie2 (http://bowtie-bio.sourceforge.net/bowtie2/index.shtml) and were deduplicated using MarkDuplicates v2.16.0 of Picard Tools. To ascertain regions of chromatin accessibility and ChIP-seq target enrichment, MACS2 (https://github.com/taoliu/MACS) was used with a p-value setting of 0.001 and scored against a publicly available input sample for cell line A375 (GSM3212794). The BEDTools suite (http://bedtools.readthedocs.io) was used to create normalized read density profiles. A global peak atlas was created by first removing blacklisted regions (http://mitra.stanford.edu/kundaje/akundaje/release/blacklists/hg38-human/hg38.blacklist.bed.gz) then merging all peaks within 500 bp for ChIP-seq or taking 500 bp windows around peak summits for ATAC and counting reads with version 1.6.1 of featureCounts (http://subread.sourceforge.net). Data were normalized by sequencing depth (to ten million unique alignments) and DESeq2 was used to calculate differential enrichment for all pairwise contrasts. Peak-gene associations were created by assigning all intragenic peaks to that gene, and otherwise using linear genomic distance to transcription start site. Peak intersections were calculated using bedtools v2.29.1 and intersectBed with 1 bp overlap. Gene set enrichment analysis (GSEA, http://software.broadinstitute.org/gsea) was performed with the pre-ranked option and default parameters, where each gene was assigned the single peak with the largest (in magnitude) log2 fold change associated with it. Motif signatures were obtained using Homer v4.5 (http://homer.ucsd.edu). Composite plots were created using deepTools v3.3.0 by running computeMatrix and plotHeatmap on normalized bigwigs with average signal sampled in 25 bp windows and flanking region defined by the surrounding 2 kb.

### Transcriptome data analysis

RNA sequencing reads were 3’ trimmed for base quality 15 and adapter sequences using version 0.4.5 of TrimGalore (https://www.bioinformatics.babraham.ac.uk/projects/trim_galore), and then aligned to human assembly hg38 with STAR v2.4 using default parameters. Data quality and transcript coverage were assessed using the Picard tool CollectRNASeqMetrics (http://broadinstitute.github.io/picard/). Read count tables were generated with HTSeq v0.9.1. Normalization and expression dynamics were evaluated with DESeq2 using the default parameters and outliers were assessed by sample grouping in principal component analysis. Gene set enrichment analysis (GSEA, http://software.broadinstitute.org/gsea) was run against MSigDB v6 using the pre-ranked option and log2 fold change for pairwise comparisons. Gene ontology biological process analysis (GOBP) was run against MSigDB v7, GO:Biological Processes, using the pre-ranked option and log2 fold change for pairwise comparisons.

### Plasmid construction for zebrafish CRISPR single guide RNA (sgRNA)

CRISPR sgRNAs of interest (sequences in **Ext. Table 11**) were cross-referenced between the top rankings from CHOPCHOP (https://chopchop.cbu.uib.no/) and integrated DNA technologies predesigned guide RNA (https://www.idtdna.com/). All oligo-synthesized sgRNAs (Forward: 5’- TTCG-(N)_19-20_-3’; Reverse: 5’-AAAC-(N)_19-20_-3’) were designed and cloned in a zebrafish U6 promoter-driven vector (Addgene#64245) at BsmBI restriction sites according to the previous study (Jao et al., 2013).

### TEAZ knockout validation using CRISPR sequencing analysis (CRISPR-seq)

Tumors generated by electroporation were dissected out from randomly selected fish. Mince the tumor tissue into small sizes with a razor and transfer the pieces immediately to a 15 ml conical tube containing tissue digestion buffer. After tissue dissociation, suspension cells were filtered through a 70 mm strainer to remove tissue debris followed by counterstaining with DAPI. After FACS, tdTomato^+^ and DAPI^-^ melanomas were lysed for phenol-chloroform based whole genome extraction. To validate gene editing using CRISPR-seq, primer pairs (**Ext. Table 11**) were set to flank 200-280 bp of the sequence that should encompass mutation sites within 100 bp from the beginning or end of it. The sequence was amplified from 50-100 ng of genomic DNA using Phusion High-Fidelity DNA Polymerase master mix PCR protocol (NEB #M0530). Gel purified PCR amplicons were sent to Integrated Genomic Operation (IGO) core facility at MSKCC for NovaSeq paired-end sequencing analysis. The results of the amplicon were aligned to zebrafish assembly GRCz10 and percent of indels at the target site was quantified using R version 4.0.5 and CrispRVariants 1.18.0 as previously described (Lindsay et al., 2016).

### Small interfering RNA (siRNA) transfection

Non-targeting Control human siRNA (#D-001810-01-05), HADHA human siRNA SMARTPool (#L-009470-00-0005), ACLY human siRNA SMARTPool (#L-004915-00-0005) were purchased from Horizon Discovery. 5.0 × 10^5^ A375 cells were plated in 6-well plates at 60% confluence with DMEM medium containing 10% FBS at a 37°C, 5% CO_2_ incubator prior to transfection. siRNAs were mixed with DharmaFECT Duo Transfection Reagent (Horizon Discovery #T-2010-03) in Opti-MEM to make 50 nM of siRNA in each well. Fresh media was added the next day followed by another 48-hr incubating for further analysis.

### Short hairpin RNA (shRNA) lentiviral transduction

To generate lentiviral particles for each shRNA, shRNA vectors were mixed with the packaging plasmid (psPAX2) and the viral envelope plasmid (pMD2.G) in a 4:2:1 ratio for transfection to 293T cells. The supernatant containing lentiviral particles was harvested at 72-hr post-transfection. The viral supernatant was applied to the target cells for a 48-hr incubation followed by puromycin selection for one week before setting up experiments.

### Reverse transcription quantitative PCR (RT-qPCR)

1.0 x 10^6^ cells were collected from 6-well plates, washed in 1x PBS, then pelleted down, followed by RNA isolation as directed by the manufacturer (Zymo Research #R1055). cDNA was then synthesized from 300-500 ng of RNA per sample using Invitrogen SuperScript IV Reverse Transcriptase (Thermo Fisher Scientific #18090050). In quantitative PCR, 1:10 ratio of cDNA was prepared as initial concentration of template in power SYBR Green PCR Master Mix for a triplicate (Thermo Fisher Scientific #4368708). Gene expression levels after DNA amplification in Bio-Rad CFX384 Real-Time detection system were calculated and normalized to internal references (GAPDH for A375 cells or hatn10 for fish melanomas) using ΔCt values.

### 3D spheroid cell invasion assay

This assay was performed based on the manufacturer’s instruction (Cultrex #3500-096). Briefly, 1.5 × 10^3^ of YUMM3.3 cells were resuspended in 50 µl of medium and were seeded in ultra-low binding, U-shaped-bottom 96 microplates followed by centrifugation to form a cluster at the center-bottom of each well. After a 72-hr incubation to allow spheroid formation, 50 µl of invasion matrix (Cultrex #3500-096-03) was gently added to embed the spheroid. After a 1-hr incubation for gel solidification, 100 µl of culture medium with vehicle control or octanoate was added to make the final 0.5 mM concentration in each well. Each spheroid was imaged by Zeiss AxioZoom system with a brightfield channel daily from day 1 to day 7 post-octanoate administration. Uniformed thresholds were applied to all images to depict the ROI of the compact core and of the total surface area through ImageJ. The manual segmentation analysis between the above-described ROIs reflects the degree of invasion defined as the invaded area in this study (Vinci et al., 2012).

### Chromatin accessibility by micrococcal nuclease (MNase) digestion

2.0 x 10^6^ A375 cells in the 100 mm dish were fixed with 4% paraformaldehyde for 10 mins followed by a 2-min incubation with 125 mM glycine after treatment with vehicle control (methanol) and 2 mM of octanoate for 20 hrs. Collected cells were resuspended in 1x PBS with 0.1% Triton X-100 (PBS-T) to a final concentration of 5.0 x 10^3^ cells/ml. 1.0 x 10^5^ cells were digested using 0.1, 10, 20, 40, 80 units of MNase (Thermo Fisher Scientific #88216) in a final volume of 50 µl of PBS-T with 5 mM CaCl_2_ at 37°C, 400 rpm agitation for 7 mins. Add 1.5 µl of 0.5M EDTA to stop the reaction. DNA purification was then performed by phenol-chloroform extraction followed by 1.2% agarose gel electrophoresis in 1x TAE buffer.

### Detection of AP-1 activity by luciferase assay

5.0 x 10^4^ cells of A375 in 24 well plates were transiently co-transfected with pRL-SV40P carrying Renilla (30 ng) as an internal control and 3xAP1pGL3 (300 ng) or pGL3 (300 ng) in a 1:10 ratio. 3xAP1pGL3 was a gift from Alexander Dent (Addgene plasmid #40342). Renilla luciferase plasmid (pRL-SV40) and Control luciferase plasmid (pGL3) were obtained from Promega (#E2231 and #E1751, respectively). 48-hr post-transfection, transfectants were treated with either LPS (Sigma-Aldrich #L2630) or octanoate. After 20-hr incubation, levels of Firefly and Renilla luminescence in cell lysates were measured by Synergy H1 microplate reader according to Promega’s Dual-Glo kit protocol (Promega #E2920). Fold changes of AP-1 activity were presented by the ratio of normalized Firefly/Renilla to control wells.

### Identification of ^13^C-labeled residues on histones H3 and H4 by liquid chromatography-mass spectrometry (LC-MS)

A375 cells in 100 mm dishes were challenged by 2 mM of octanoate-1-^13^C (Sigma Aldrich #296457) for 6 or 20 hrs. After treatment, cells were harvested and pelleted down for acidic protein extraction as previously described (Sidoli et al., 2016). To validate a successful histone precipitation, reconstituted protein extracts were resolved in pre-cast 4-15% gradient SDS-PAGE gels and sent in liquid solution for LC-MS analysis. The following steps for general sample preparation, tryptic digestion, microtip C18 desalting of peptides, LC-MS analysis were continued and performed by Weill Cornell Medicine (WCM) Proteomics and Metabolomics Core Facility. Histones were digested in solution with trypsin at 37°C, O/N. The digests were vacuum centrifuged to near dryness and desalted by micro-C18 column before subjected to LC-MS/MS analysis on a Thermo Fisher Scientific EASY-nLC 1000 coupled on-line to a Fusion Lumos mass spectrometer (Thermo Fisher Scientific). Buffer A (0.1% FA in water) and buffer B (0.1% FA in ACN) were used as mobile phases for gradient separation. A 75 µm x 15 cm chromatography column (ReproSil-Pur C18-AQ, 3 µm, Dr. Maisch GmbH, German) packed in-house was used for peptide separation. Peptides were separated with a gradient of 3–30% buffer B over 20 mins, 30-80% B over 10 mins at a flow rate of 300 nL/min. The Fusion Lumos mass spectrometer was operated in data dependent mode. Full MS scans were acquired in the Orbitrap mass analyzer over a range of 300-1600 m/z with resolution 70,000 at m/z 200. The top 15 most abundant precursors with charge states between 2 and 5 were selected with an isolation window of 1.4 Thomsons by the quadrupole and fragmented by higher-energy collisional dissociation with normalized collision energy of 35. MS/MS scans were acquired in the Orbitrap mass analyzer with resolution 15,000 at m/z 200. The automatic gain control target value was 1e6 for full scans and 5e4 for MS/MS scans respectively, and the maximum ion injection time is 60 ms for both.

The raw files were processed using the MaxQuant computational proteomics platform version 1.5.5.1 (Max Planck Institute, Munich, Germany) for protein identification. The fragmentation spectra were used to search the UniProt yeast protein database (downloaded on 10/18/2019). Oxidation of methionine, protein N-terminal acetylation, lysine acetylation, and lysine propionylation were used as variable modifications for database searching. The precursor and fragment mass tolerances were set to 7 and 20 ppm, respectively. Both peptide and protein identifications were filtered at 1% false discovery rate based on decoy search using a database with the protein sequences reversed. The result was listed in **Ext. Table 3**.

### Statistics

All the data presented with statistics were generated in GraphPad Prism 9.3.1 and conducted by at least three biological independent replicates unless otherwise mentioned in the figure legends.

## Extended Figure Legends

**Extended Figure 1.**
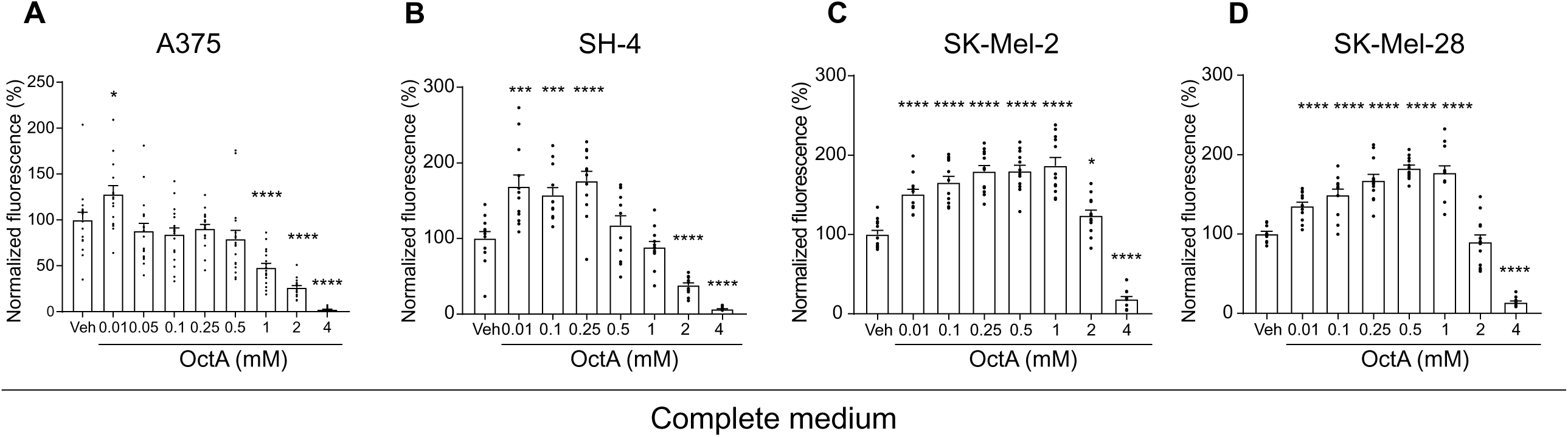
mcFA supplementation promotes cell proliferation. Examining the effect of octanoate on cell growth using human melanoma cell lines: A375 (**A**); SH-4 (**B**); SK-Mel-2 (**C**) and SK-Mel-28 (**D**). 1,000 cells were seeded in each well of 96 well plates overnight. Aspirate medium and culture cells in complete DMEM with the addition of indicated concentrations of octanoate for 5 days. Cell proliferation was then determined by CyQUANT cell proliferation assay. Mean ± SEM, n=3, unpaired t-Test. *, p<0.05; **, p<0.01; ***, p<0.005; ****, p<0.001

**Extended Figure 2.**
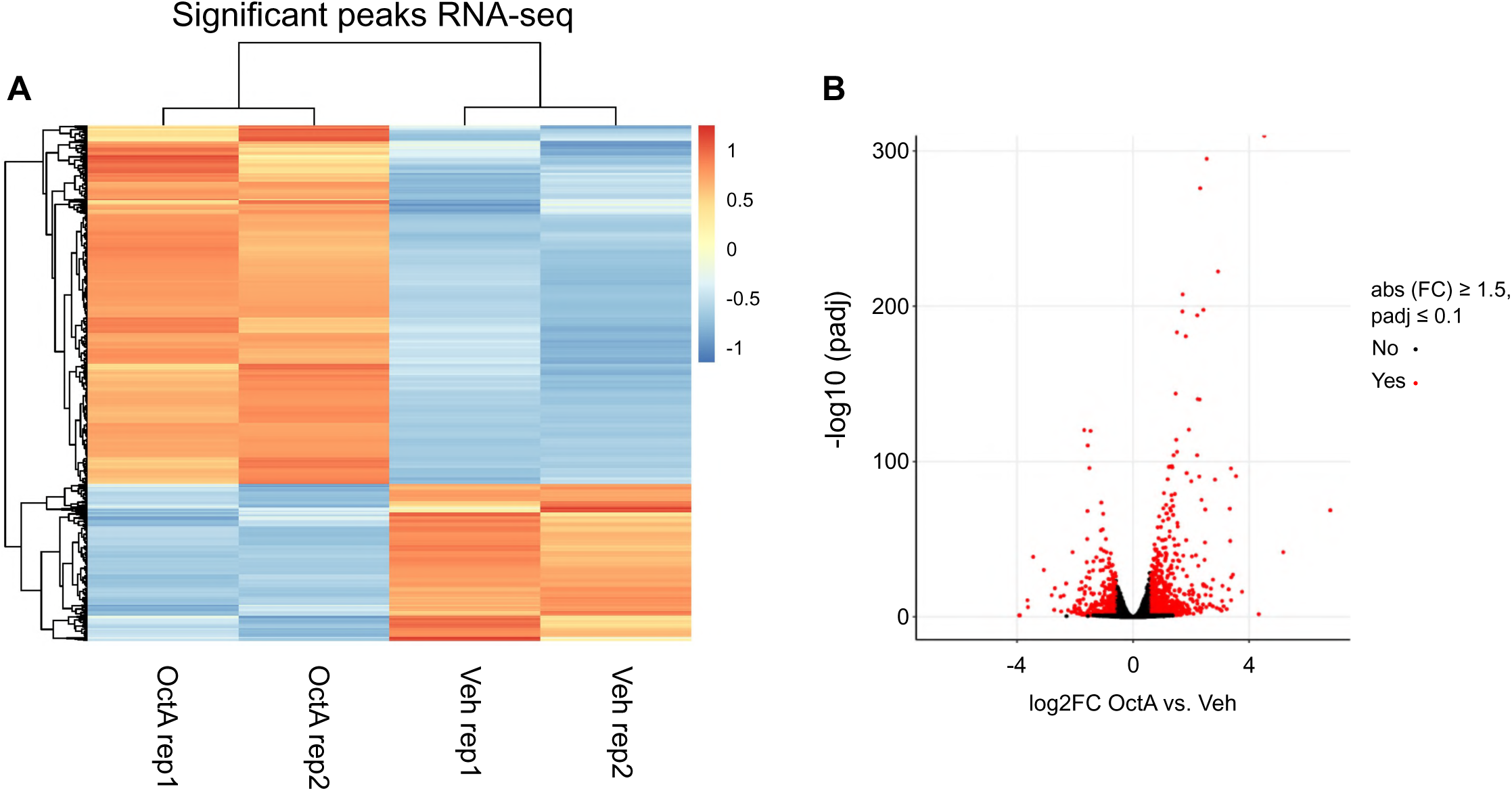
Differential expression gene analysis of bulk RNA-seq in human A375 cells with 922 significantly upregulated and 403 downregulated peaks shown by the heatmap (**A**) and the volcano plot (**B**).

**Extended Figure 3.**
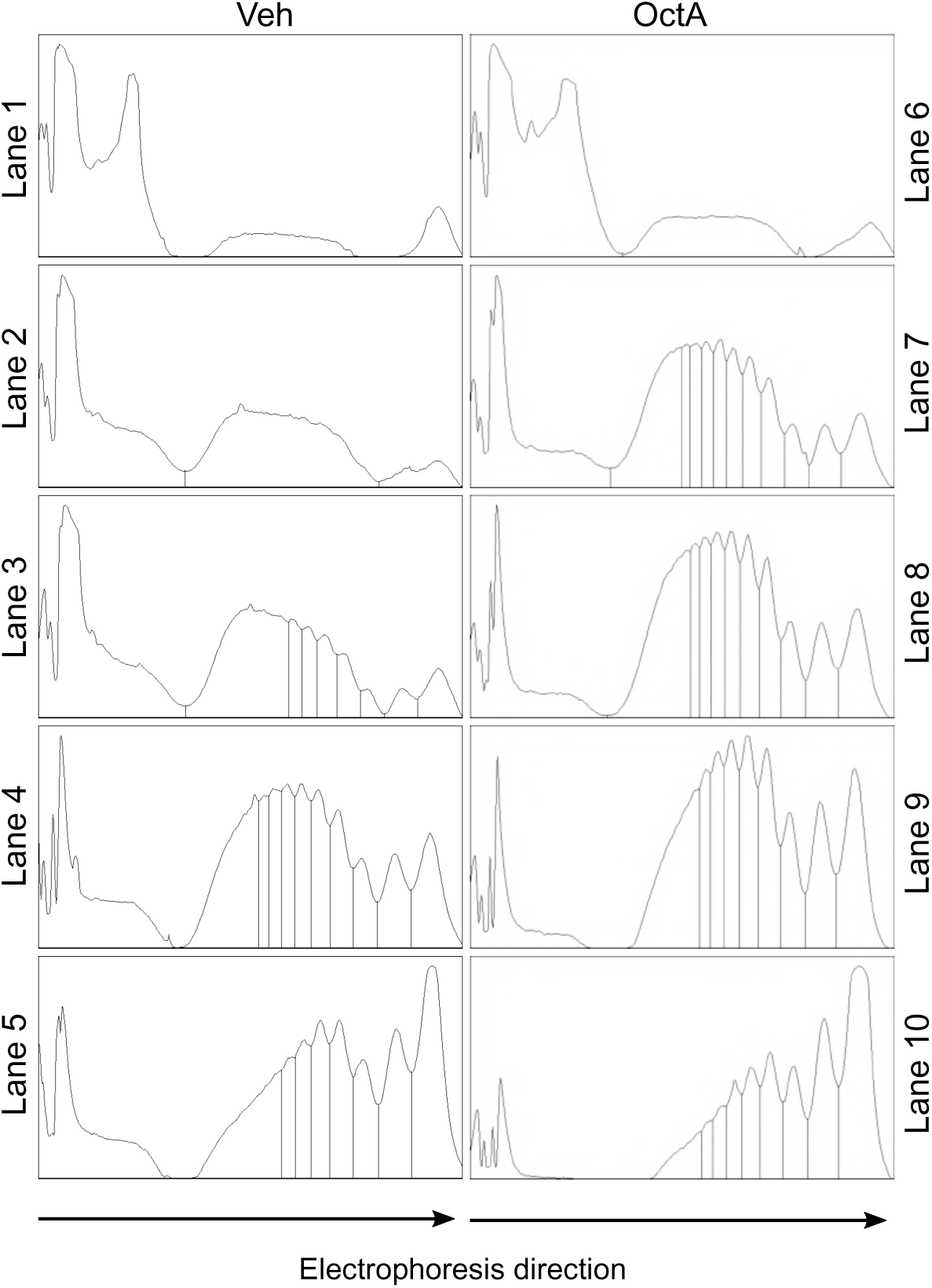
Quantification of DNA fragments in the agarose gel shown in Fig.2D. Peak distribution plots made by ImageJ represent levels of DNA fragments in each lane migrating along with the direction of electrophoresis (Left to right: “-” to “+”).

**Extended Figure 4.**
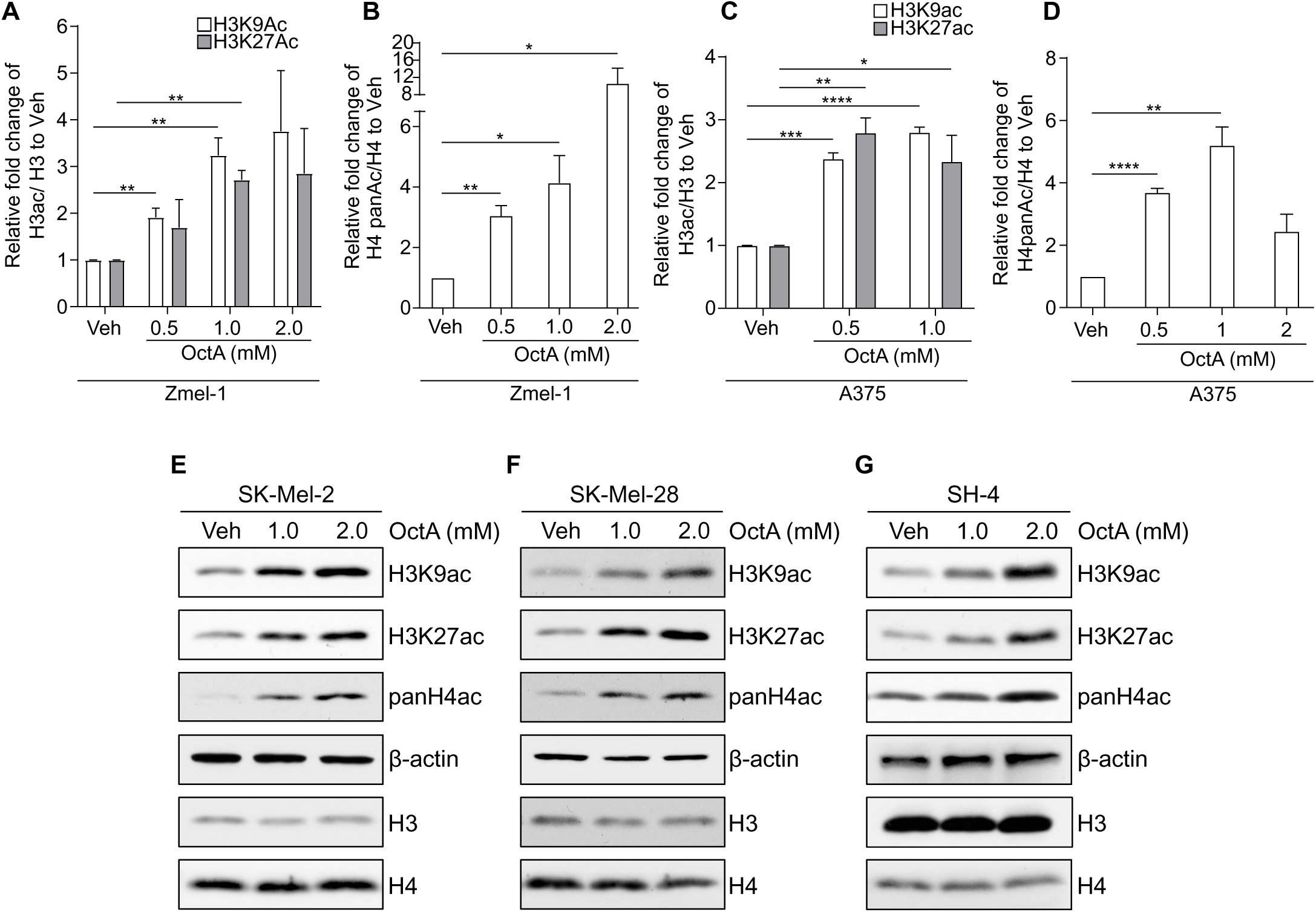
Quantification of histone acetylation in Zmel-1 (**A**, **B**) and A375 (**C**, **D**) shown in **Figs. 3A-B**. Each panel represents relative fold changes of indicated acetylated histone levels that were normalized by basal histone H3 or H4 to vehicle control. Mean ± SEM, n=3, unpaired t-Test, *, p<0.05; **, p<0.01; ***, p<0.005; ****, p<0.001. Representative of histone H3 and H4 acetylation abundance in human melanoma cell lines, SK-Mel-2 (**E**), SK-Mel-28 (**F**), and SH-4 (**G**) performed by Western blot analysis. n=3.

**Extended Figure 5.**
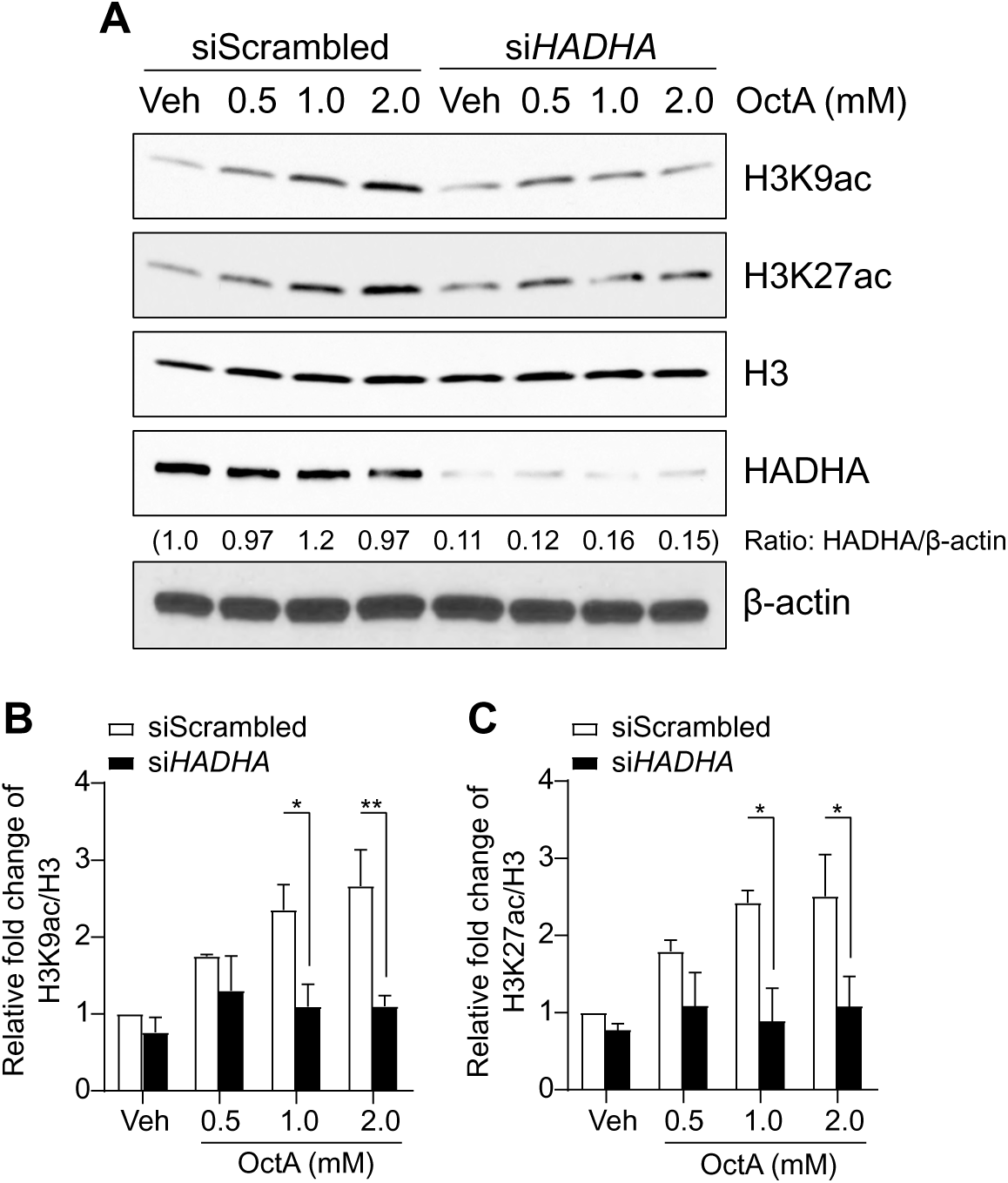
Examining the effects of HADHA on histone H3 acetylation at lysine 9 and 27 residues. Scrambled control and HADHA-silenced A375 cells were challenged by octanoate for 20 hours. Representative images of Western blot and ratios of HADHA protein were shown in (**A**). Relative levels of H3K9ac and H3K27ac after normalization to H3 were shown in (**B**) and (**C**), respectively. Mean ± SEM, n=3, 2-way ANOVA, *, p<0.05; **, p<0.01.

**Extended Figure 6.**
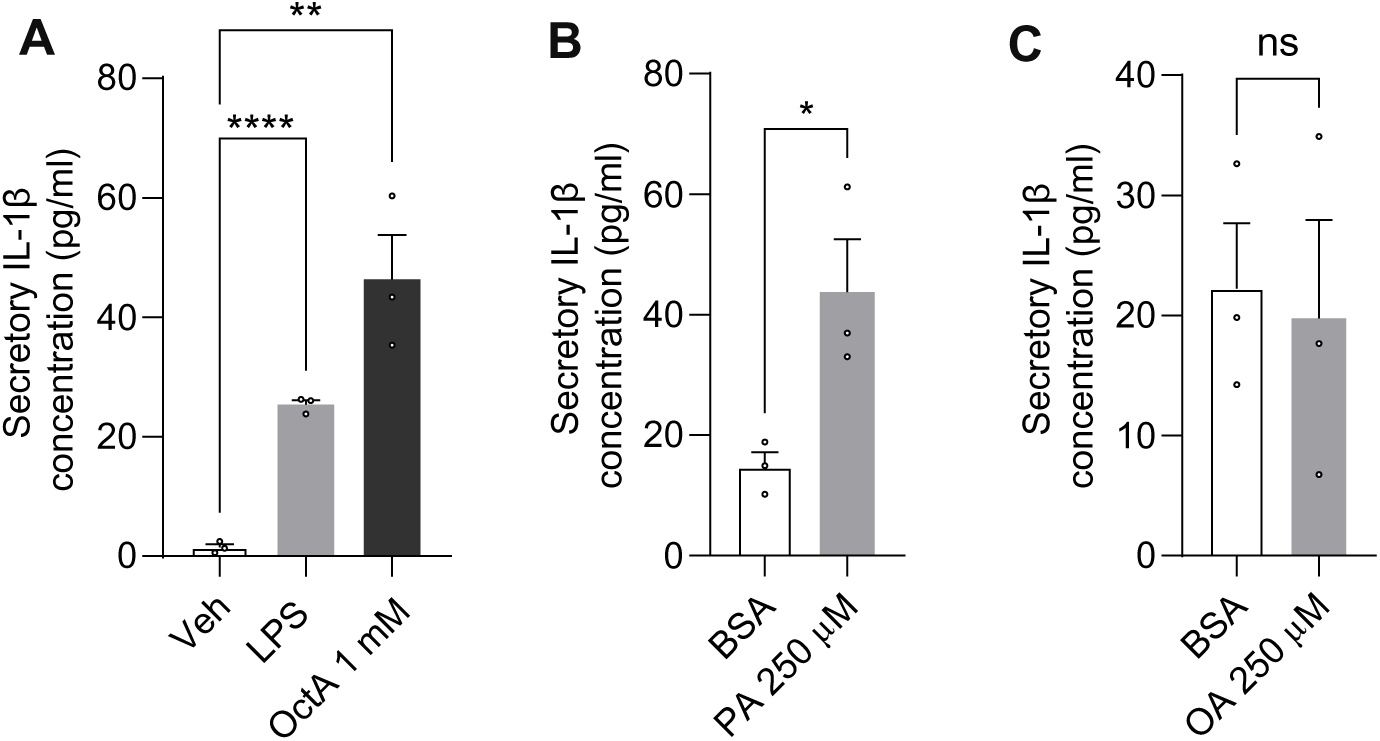
Levels of secretory IL-1β in response to FAs by ELISA analysis. A375 cells were stimulated by LPS (1 µg/ml) or octanoate (**A**) or palmitate (**B**) or oleate (**C**) for 72 hrs. Collected media were applied to the ELISA assay according to the manufacturer’s instruction. Mean ± SEM, n=3, unpaired t-Test, **, p<0.01; ****, p<0.001.

**Extended Figure 7.**
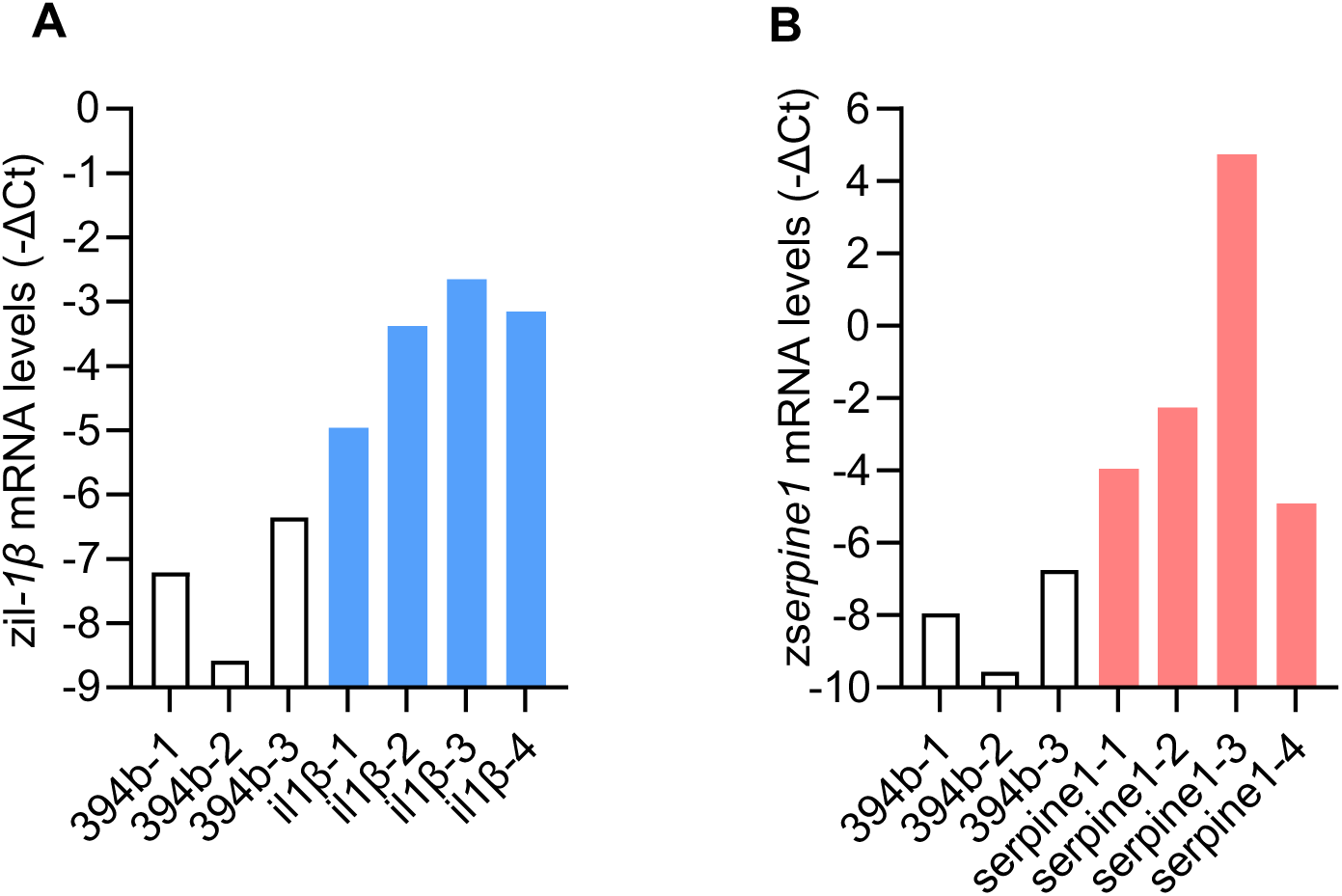
Validation of exogenous cytokine expression in fish melanoma. The melanoma tissue was dissected from randomly selected fish after 8-wpe. Tumor samples after one time wash in 1x PBS were directly used for RNA extraction followed by cDNA synthesis. The mRNA level of *il-1β* (**A**) or *serpine1* (**B**) in representative fish was shown by (-ΔCt) via qPCR analysis.

**Extended Figure 8.**
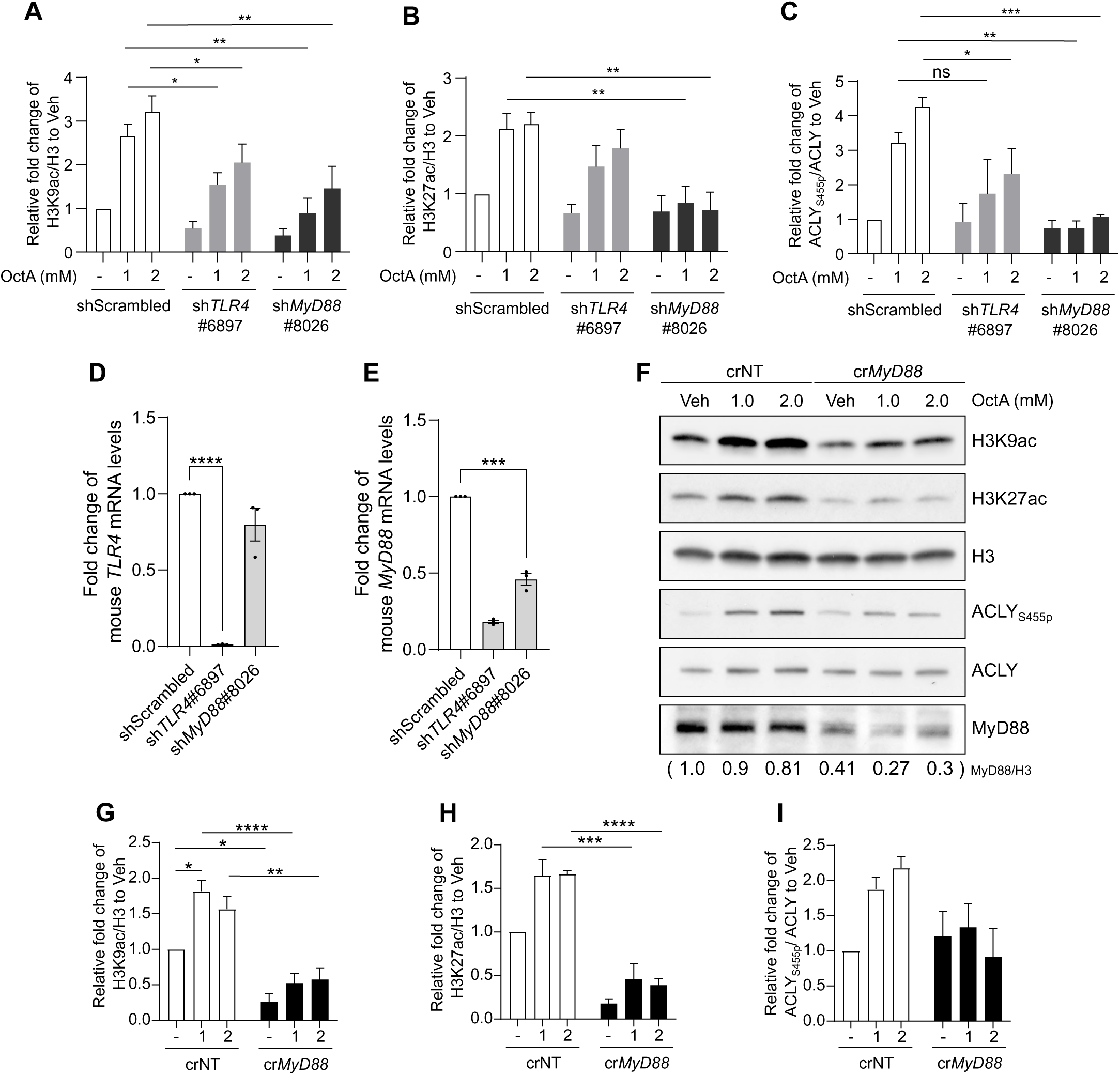
Assessing the effects of TLR4 and MyD88 on octanoate-induced H3 acetylation and ACLY phosphorylation in human A375 cells. (**A**-**C**) Quantification of H3K9ac (**A**), H3K27ac (**B**) and ACLY_S455p_ (**C**) in Fig.5D. Mean ± SEM, n=3, 2-way ANOVA, *, p<0.05; **, p<0.01; ***, p<0.005. Knockdown of human *TLR4* (**D**) and *MyD88* (**E**) was shown by qPCR. Mean ± SEM, n=3, unpaired t-Test, *, p<0.005; ****, p<0.001. To generate CRISPR-mediated *MyD88^KO^* cells, A375 cells were transfected with the double nickase plasmid for control (Santa Cruz #sc-437281) or *MyD88* (Santa Cruz #SC-417166-NIC-2) using DharmaFECT Duo Transfection Reagent. After transfection, cells went through puromycin selection for a week and were incubated in fresh medium for 2 weeks or until starting expansion otherwise. Pool clones of knockout cells were set for 3 biological experiments and the Western blot result was shown in (**F**) with quantification of H3ac (**G**, **H**) and ACLY phosphorylation (**I**). Mean ± SEM, n=3, 2-way ANOVA, *, p<0.05; **, p<0.01; ***, p<0.005; ****, p<0.001.

**Extended Figure 9.**
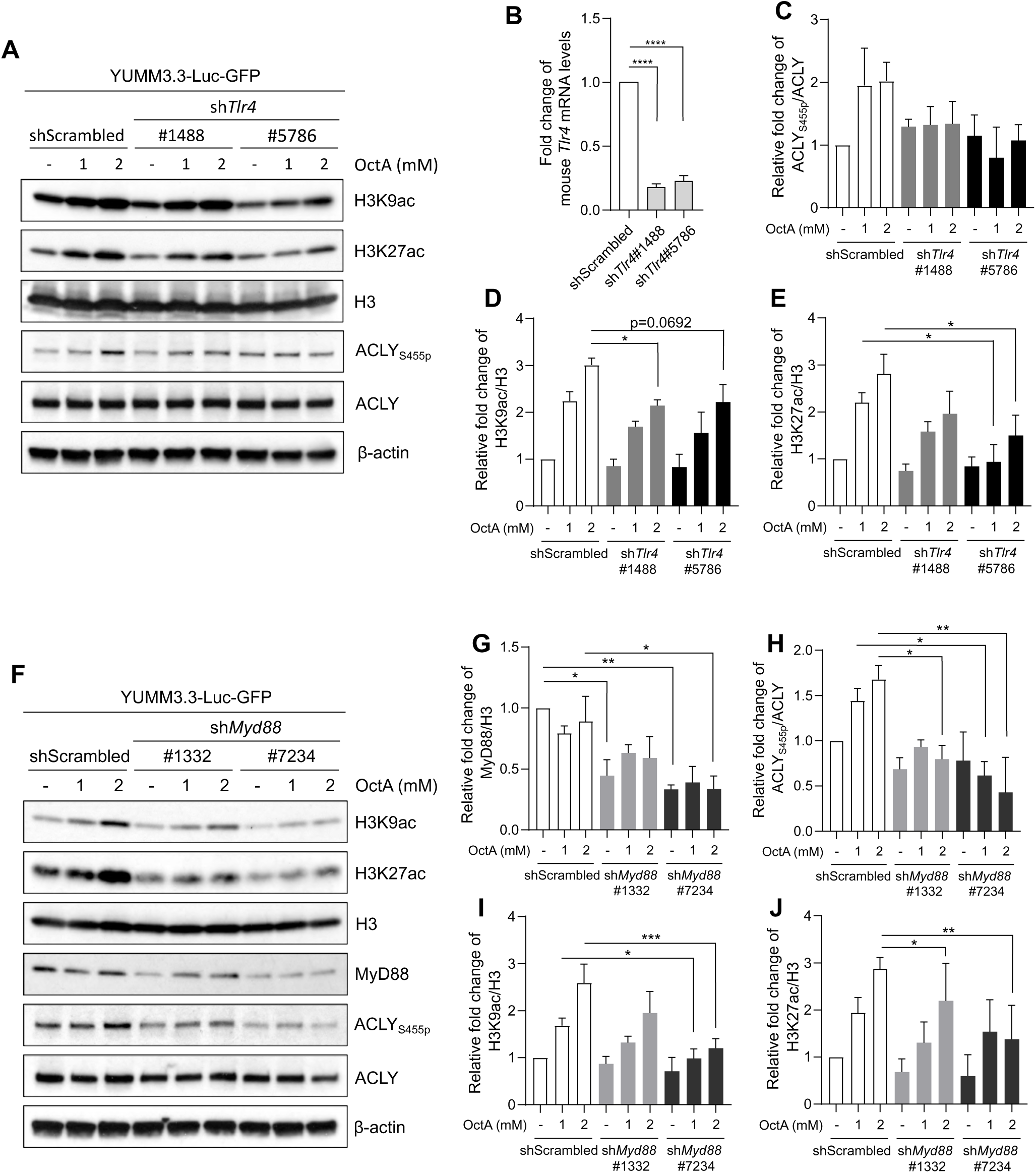
Western blot analysis of H3 acetylation and ACLY phosphorylation in octanoate-treated mouse YUMM3.3 cells lacking Tlr4 (**A**) or Myd88 (**F**). Two shRNAs for each gene were individually used in this experiment. Knockdown of *Tlr4* (**B**) or *Myd88* (**G**) was examined by qPCR analysis. Mean ± SEM, n=3, unpaired t-Test, *, p<0.05; **, p<0.01; ****, p<0.001. Quantification of H3ac and ACLY phosphorylation shown in (**C**-**E**) and (**H**-**J**) as indicated. Mean ± SEM, n=3, 2-way ANOVA, *, p<0.05; **, p<0.01; ***, p<0.005.

**Extended Figure 10.**
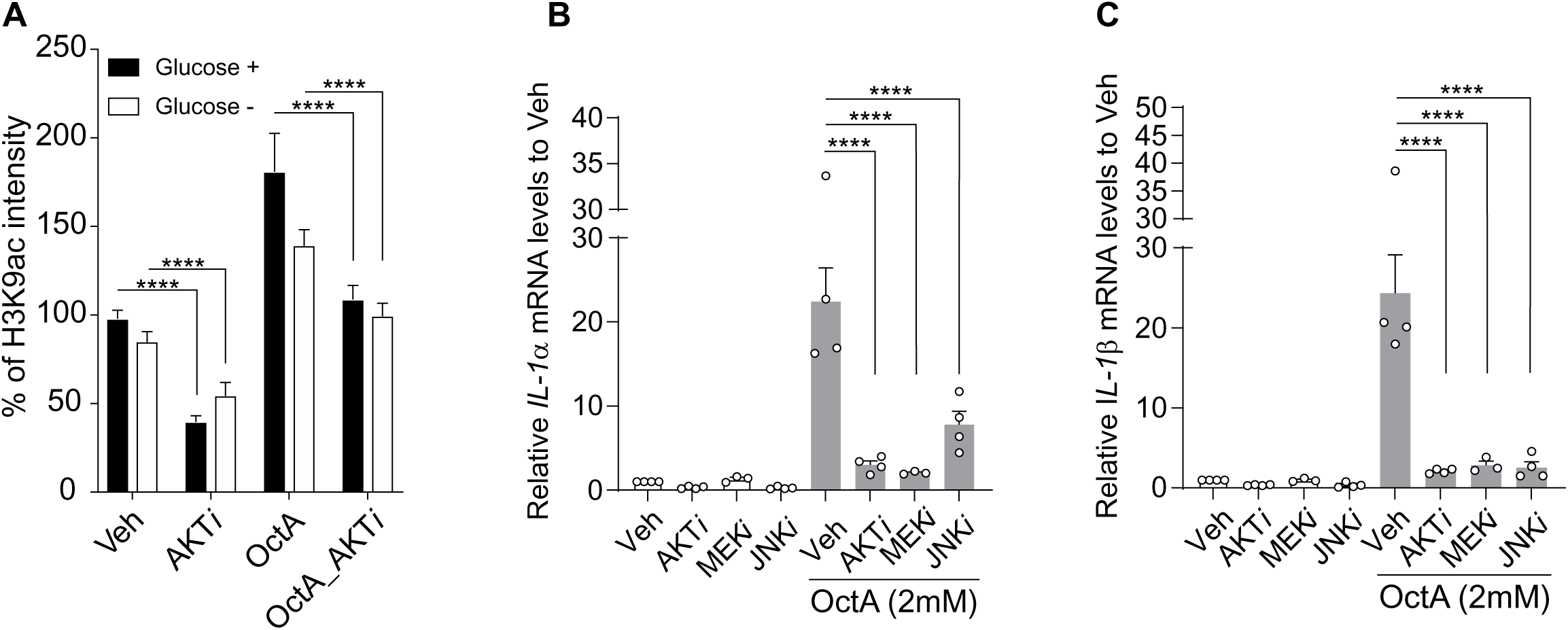
Investigating the effects of pharmacological inhibition of AKT, JNK, and MEK kinases on H3K9 acetylation and *IL-1* expression. (**A**) A375 cells in 96-well plates of were pretreated for 1 hr with 10 µM AKT inhibitor GSK690693 (Selleck #S1113) and then co-treated for 20 hours with 2 mM octanoate in complete or glucose-free DMEM. Following immunofluorescent staining for H3K9ac, images were captured using IN Cell Analyzer model 4002. Mean ± SEM, n=2, unpaired t-test, ****, p<0.001. The A375 cells were pretreated with 10 µM AKT inhibitor or 10 µM of JNK inhibitor SP600125 (Selleck #S1460) or 10 µM of MEK inhibitor PD0325901 (Sigma-Aldrich #PZ0162) followed by the addition of 2 mM of octanoate to 6-well plates. *IL-1α* (**B**) and *IL-1β* (**C**) mRNA levels were measured by qPCR analysis. Mean ± SEM, n=3, unpaired t-test, ****, p<0.001.

**Extended Figure 11.**
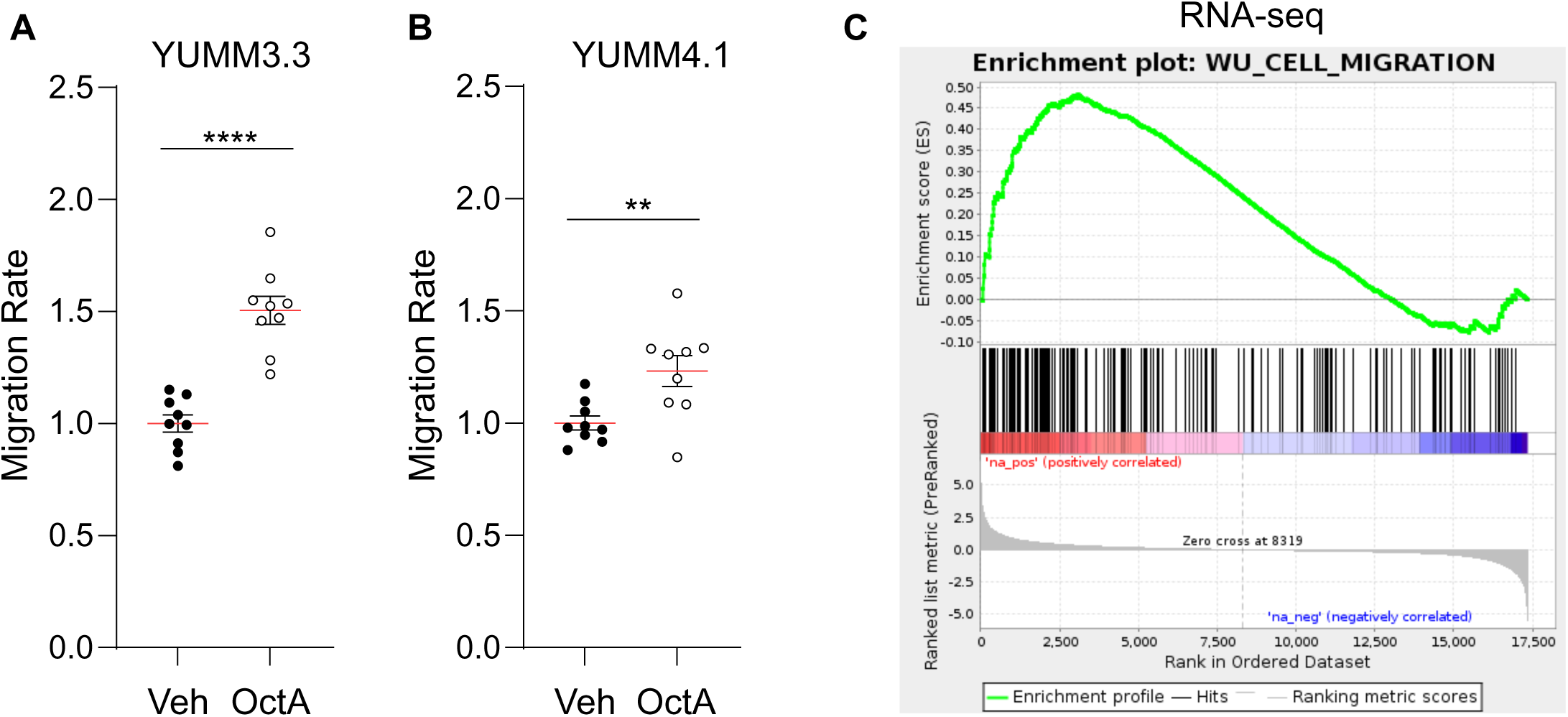
Octanoate enhances melanoma migration. YUMM3.3 cells were primed with vehicle or 1 mM octanoate for 5 days before being tested using the Cultrex invasion assay (R&D systems #3457-096-K). Briefly, the 96-well top invasion chamber was coated with 0.5 x collagen I solution at 37°C, overnight. 5.0 x 10^4^ cells in 50 µl of medium containing 0.5% FBS were added to the top chamber and the bottom chamber was filled with 150 µl of complete DMEM. Assay chambers were incubated at 37°C for 18 hours followed by the rest of the manufacturer’s instructions. The migrating capability of YUMM3.3 (**A**) or YUMM4.1 (**B**) was shown by 3 biological replicates. Each dot represents one technical replicate. 3 technical replicates were set for one independent experiment. Mean ± SEM, unpaired t-Test, **, p<0.01; ****, p<0.001. (**C**) Enrichment of cell migration gene signature (OctA vs. Veh) in RNA-seq.

**Extended Figure 12.**
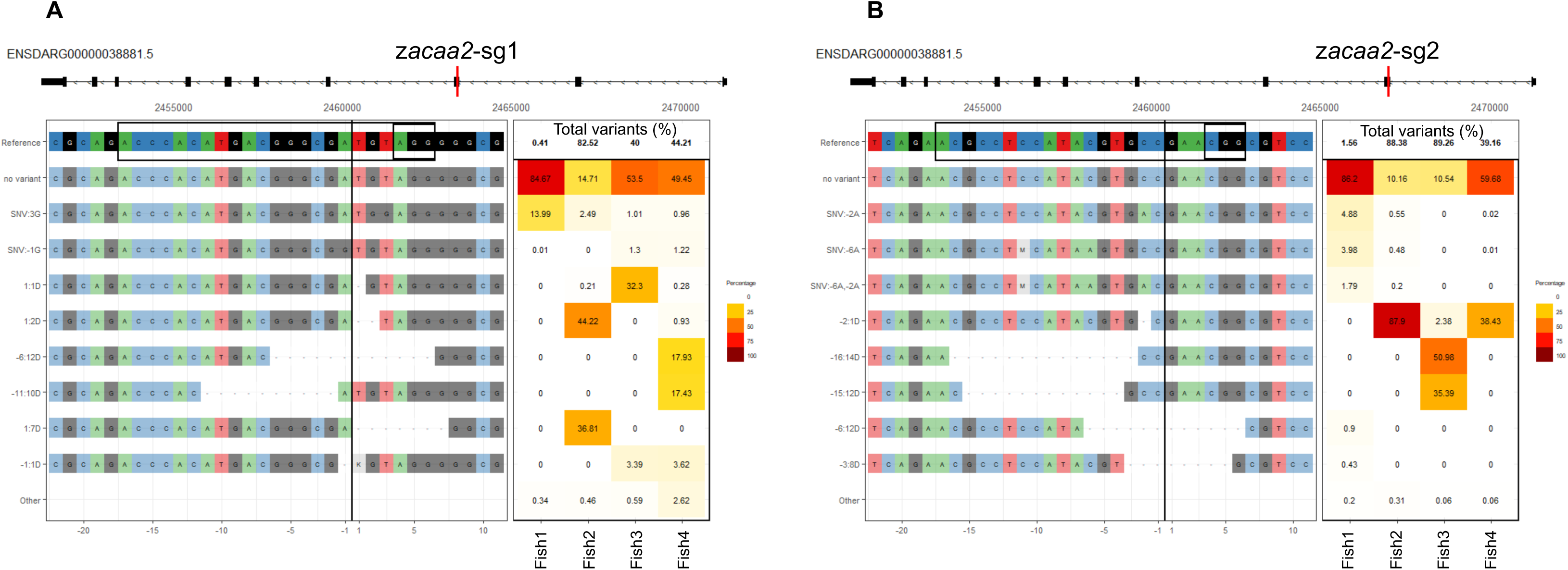
Validation of *in vivo* z*acaa2* knockout via CRISPR-seq. Genomic DNA from the same fish tumor was allocated to z*acaa2*-sg1 and z*acaa2*-sg2 PCR amplification. Variants of genetic mutations generated by CRISPR-z*acaa2*-sg1 (**A**) and CRISPR-z*acaa2*-sg2 (**B**). Genomic locus of the guide sequence was shown in red.

**Extended Figure 13.**
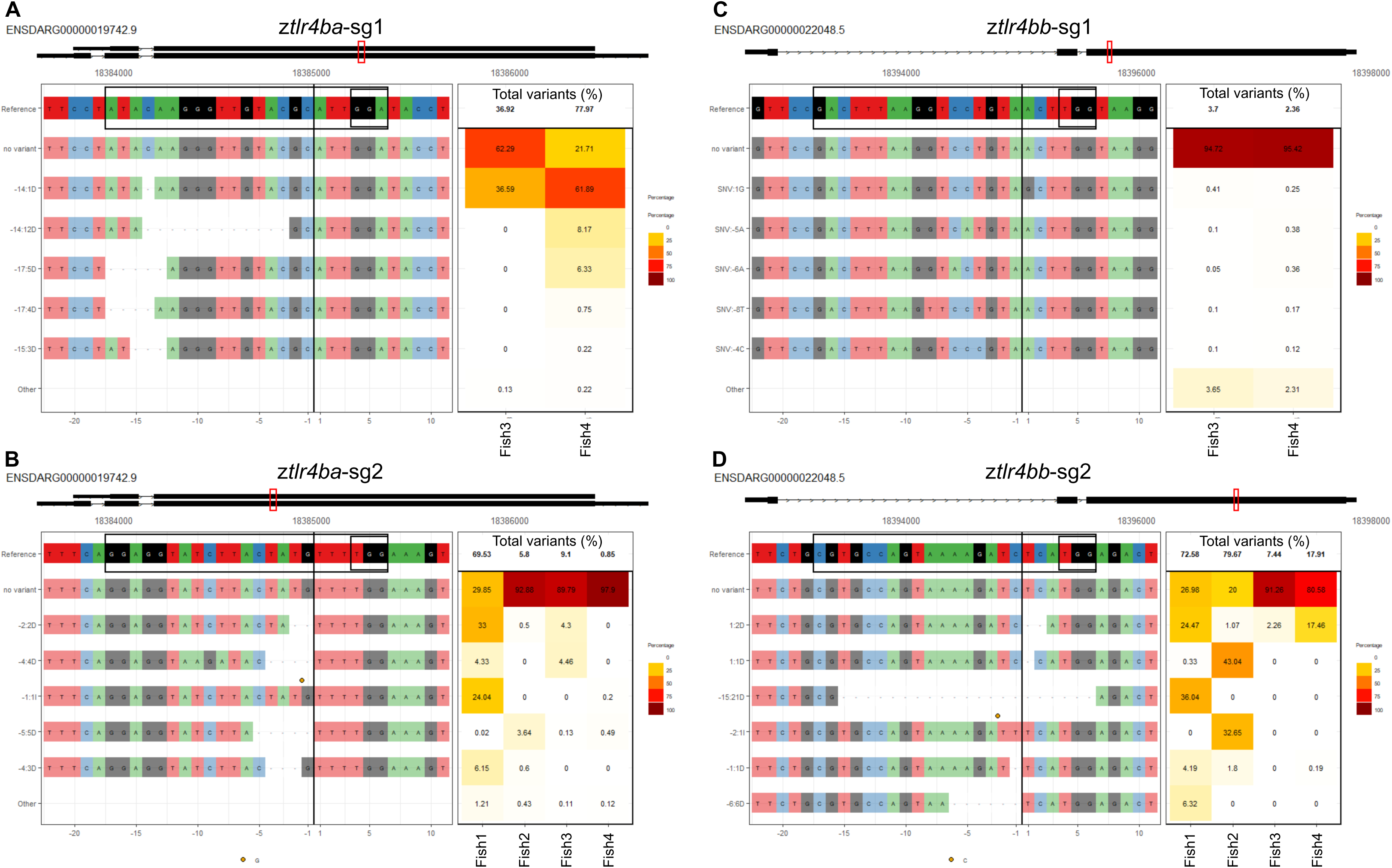
Validation of *in vivo* z*tlr4* knockout by CRISPR-seq. Genomic DNA from the same fish tumor was divided for z*tlr4ba*-sg1, z*tlr4ba*-sg2, z*tlr4bb*-sg1, and z*tlr4bb*-sg2 PCR amplification. Variants of genetic mutations generated by CRISPR-z*tlr4ba*-sg1 (**A**), CRISPR-z*tlr4ba*-sg2 (**B**), CRISPR-z*tlr4bb*-sg1 (**C**) and CRISPR-z*tlr4bb*-sg2 (**D**). Genomic locus of the guide sequence was shown in red.

**Extended Figure 14.**
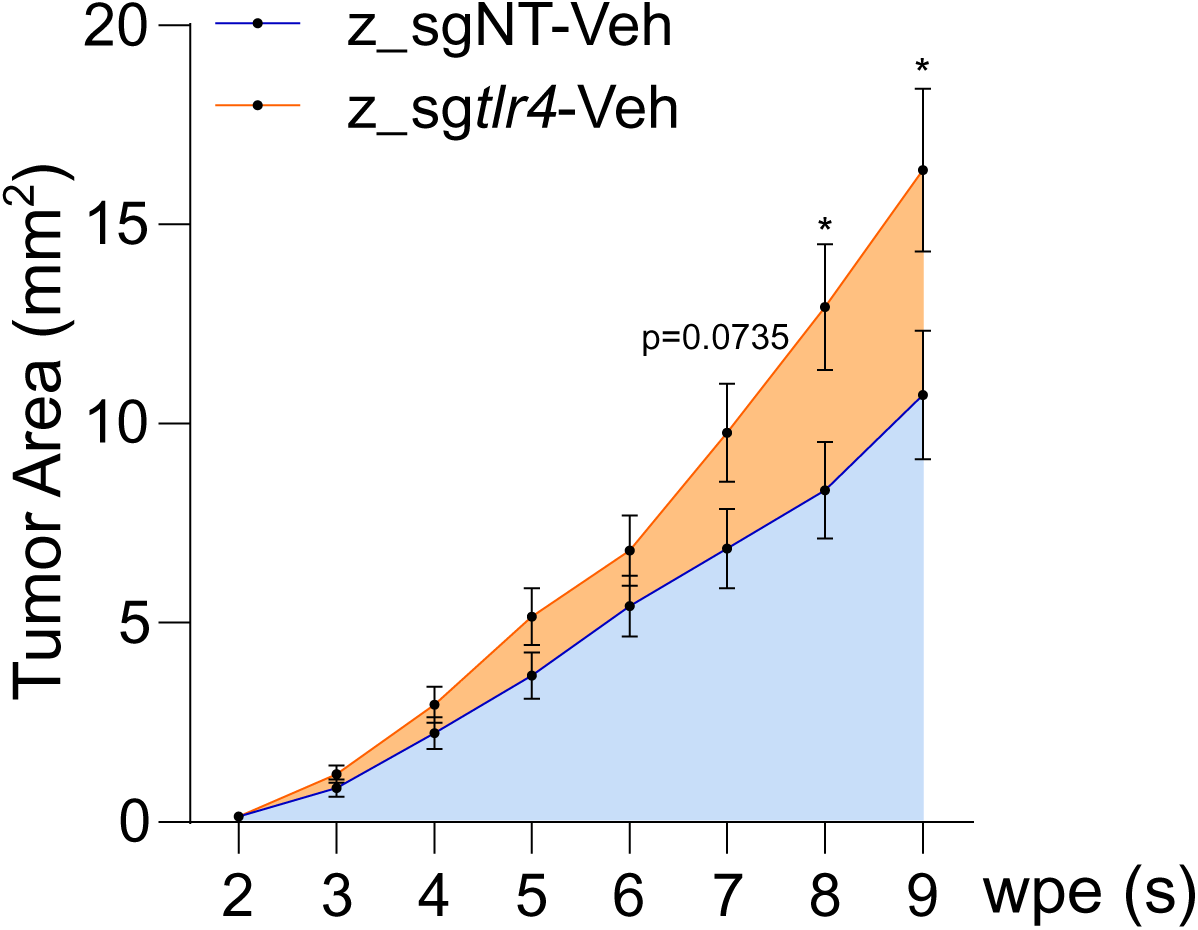
Comparison of tumor growth between non-targeting and *tlr4^KO^*TEAZ fish. The data of Veh were extracted from Fig.6F and Fig.6H. Mean ± SEM, unpaired t-Test, *, p<0.05.

**Extended Figure 15.**
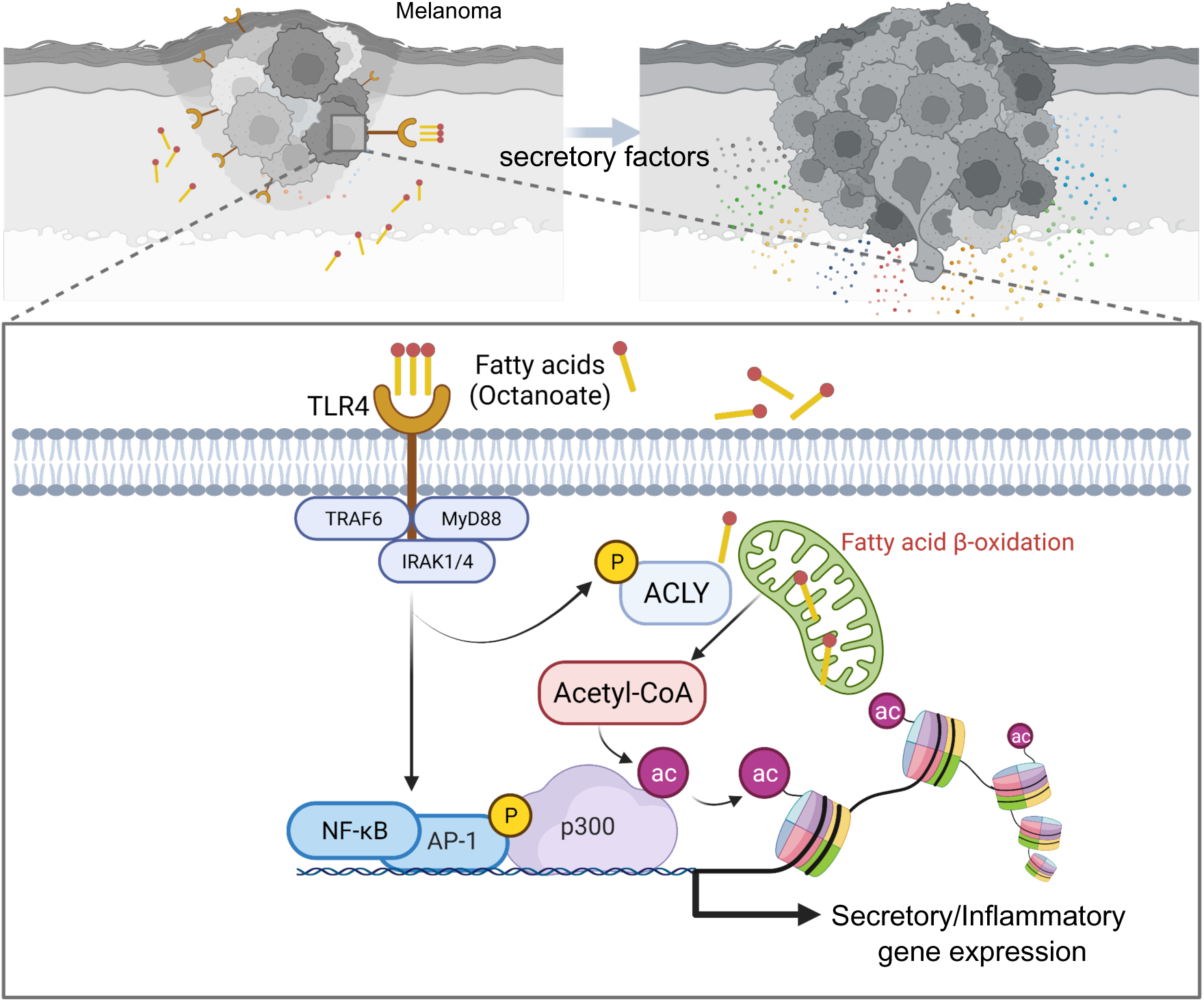
Model showing how the mcFA octanoate promotes melanoma progression via the cooperation between FA metabolism and TLR4 signaling. Made by BioRender.

## Extended Table Index

**Extended Table 1**

GSEA analysis of RNA-seq using all gene sets of MSigDB

**Extended Table 2**

GSEA analysis of RNA-seq by GOBP annotation

**Extended Table 3**

13C-labeled histone residues by LC-MS analysis

**Extended Table 4**

GSEA analysis of H3K27ac ChIP-seq using all gene sets of MSigDB

**Extended Table 5**

Common upregulated DEGs between RNA-seq and H3K27ac ChIP-seq

**Extended Table 6**

PANTHER Gene List-common DEGs between RNA-seq, ChIP-seq, and HPA secretome

**Extended Table 7**

Significant DEGs (OctA vs. Veh) of RNA-seq

**Extended Table 8**

Table of multi-plasmid mixture for TEAZ

**Extended Table 9**

Primary antibodies used in this study

**Extended Table 10**

Short hairpin RNAs (shRNAs) used in mouse and human cells.

**Extended Table 11**

Primer sets for CRISPR-seq and single guide sequence of target genes in zebrafish.

**Extended Table 12**

Primer sets for qPCR analysis.

**Extended Table 13**

Published available and filtered HPA secretome database.

**Extended Table 14**

Significant DEGs (OctA vs. Veh) of H3K27ac ChIP-seq

**Extended Table 15**

All normalized counts of RNA-seq

**Extended Table 16**

All normalized foldpv of H3K27ac ChIP-seq

## Notes

### Competing Interest Statement

The authors have declared no competing interest.

## References

Ahmed, A., Wang, J.H., and Redmond, H.P. (2013). Silencing of TLR4 increases tumor progression and lung metastasis in a murine model of breast cancer. Ann. Surg. Oncol. 20 *Suppl 3*, S389–96.

Baenke, F., Peck, B., Miess, H., and Schulze, A. (2013). Hooked on fat: the role of lipid synthesis in cancer metabolism and tumour development. Dis. Model. Mech. 6, 1353–1363.

Baker, K.J., Houston, A., and Brint, E. (2019). IL-1 Family Members in Cancer; Two Sides to Every Story. Front. Immunol. 10, 1197.

Bald, T., Quast, T., Landsberg, J., Rogava, M., Glodde, N., Lopez-Ramos, D., Kohlmeyer, J., Riesenberg, S., van den Boorn-Konijnenberg, D., Hömig-Hölzel, C., et al. (2014). Ultraviolet-radiation-induced inflammation promotes angiotropism and metastasis in melanoma. Nature 507, 109–113.

Batista-Gonzalez, A., Vidal, R., Criollo, A., and Carreño, L.J. (2019). New insights on the role of lipid metabolism in the metabolic reprogramming of macrophages. Front. Immunol. 10, 2993.

Block, M.S., Vierkant, R.A., Rambau, P.F., Winham, S.J., Wagner, P., Traficante, N., Tołoczko, A., Tiezzi, D.G., Taran, F.A., Sinn, P., et al. (2018). Myd88 and TLR4 expression in epithelial ovarian cancer. Mayo Clin. Proc. 93, 307–320.

Buenrostro, J.D., Giresi, P.G., Zaba, L.C., Chang, H.Y., and Greenleaf, W.J. (2013). Transposition of native chromatin for fast and sensitive epigenomic profiling of open chromatin, DNA-binding proteins and nucleosome position. Nat. Methods 10, 1213–1218.

Bugyei-Twum, A., Advani, A., Advani, S.L., Zhang, Y., Thai, K., Kelly, D.J., and Connelly, K.A. (2014). High glucose induces Smad activation via the transcriptional coregulator p300 and contributes to cardiac fibrosis and hypertrophy. Cardiovasc. Diabetol. 13, 89.

Burkholder, B., Huang, R.-Y., Burgess, R., Luo, S., Jones, V.S., Zhang, W., Lv, Z.-Q., Gao, C.- Y., Wang, B.-L., Zhang, Y.-M., et al. (2014). Tumor-induced perturbations of cytokines and immune cell networks. Biochim. Biophys. Acta 1845, 182–201.

Callahan, S.J., Tepan, S., Zhang, Y.M., Lindsay, H., Burger, A., Campbell, N.R., Kim, I.S., Hollmann, T.J., Studer, L., Mosimann, C., et al. (2018). Cancer modeling by Transgene Electroporation in Adult Zebrafish (TEAZ). Dis. Model. Mech. 11.

Campbell, S.L., and Wellen, K.E. (2018). Metabolic signaling to the nucleus in cancer. Mol. Cell 71, 398–408.

Carrer, A., Trefely, S., Zhao, S., Campbell, S.L., Norgard, R.J., Schultz, K.C., Sidoli, S., Parris, J.L.D., Affronti, H.C., Sivanand, S., et al. (2019). Acetyl-CoA Metabolism Supports Multistep Pancreatic Tumorigenesis. Cancer Discov. 9, 416–435.

Ceol, C.J., Houvras, Y., Jane-Valbuena, J., Bilodeau, S., Orlando, D.A., Battisti, V., Fritsch, L., Lin, W.M., Hollmann, T.J., Ferré, F., et al. (2011). The histone methyltransferase SETDB1 is recurrently amplified in melanoma and accelerates its onset. Nature 471, 513–517.

Chen, S., Feng, B., George, B., Chakrabarti, R., Chen, M., and Chakrabarti, S. (2010). Transcriptional coactivator p300 regulates glucose-induced gene expression in endothelial cells. Am. J. Physiol. Endocrinol. Metab. 298, E127–37.

Chen, X., Zhao, F., Zhang, H., Zhu, Y., Wu, K., and Tan, G. (2015). Significance of TLR4/MyD88 expression in breast cancer. Int. J. Clin. Exp. Pathol. 8, 7034–7039.

Clement, E., Lazar, I., Attané, C., Carrié, L., Dauvillier, S., Ducoux-Petit, M., Esteve, D., Menneteau, T., Moutahir, M., Le Gonidec, S., et al. (2020). Adipocyte extracellular vesicles carry enzymes and fatty acids that stimulate mitochondrial metabolism and remodeling in tumor cells. EMBO J. 39, e102525.

Coutzac, C., Jouniaux, J.-M., Paci, A., Schmidt, J., Mallardo, D., Seck, A., Asvatourian, V., Cassard, L., Saulnier, P., Lacroix, L., et al. (2020). Systemic short chain fatty acids limit antitumor effect of CTLA-4 blockade in hosts with cancer. Nat. Commun. 11, 2168.

Currie, E., Schulze, A., Zechner, R., Walther, T.C., and Farese, R.V. (2013). Cellular fatty acid metabolism and cancer. Cell Metab. 18, 153–161.

Djouadi, F., Habarou, F., Le Bachelier, C., Ferdinandusse, S., Schlemmer, D., Benoist, J.F., Boutron, A., Andresen, B.S., Visser, G., de Lonlay, P., et al. (2016). Mitochondrial trifunctional protein deficiency in human cultured fibroblasts: effects of bezafibrate. J. Inherit. Metab. Dis. 39, 47–58.

Erridge, C., and Samani, N.J. (2009). Saturated fatty acids do not directly stimulate Toll-like receptor signaling. Arterioscler. Thromb. Vasc. Biol. 29, 1944–1949.

Fan, L., Lin, Q., Huang, X., Fu, D., and Huang, H. (2021). Prognostic significance of pretreatment serum free fatty acid in patients with diffuse large B-cell lymphoma in the rituximab era: a retrospective analysis. BMC Cancer 21, 1255.

Fenton, M.J., Clark, B.D., Collins, K.L., Webb, A.C., Rich, A., and Auron, P.E. (1987). Transcriptional regulation of the human prointerleukin 1 beta gene. J. Immunol. 138, 3972– 3979.

Fritsche, K.L. (2015). The science of fatty acids and inflammation. Adv. Nutr. 6, 293S–301S.

Fukata, M., Chen, A., Vamadevan, A.S., Cohen, J., Breglio, K., Krishnareddy, S., Hsu, D., Xu, R., Harpaz, N., Dannenberg, A.J., et al. (2007). Toll-like receptor-4 promotes the development of colitis-associated colorectal tumors. Gastroenterology 133, 1869–1881.

Fu, X.-Q., Liu, B., Wang, Y.-P., Li, J.-K., Zhu, P.-L., Li, T., Tse, K.-W., Chou, J.-Y., Yin, C.-L., Bai, J.-X., et al. (2020). Activation of STAT3 is a key event in TLR4 signaling-mediated melanoma progression. Cell Death Dis. 11, 246.

Gay, N.J., Symmons, M.F., Gangloff, M., and Bryant, C.E. (2014). Assembly and localization of Toll-like receptor signalling complexes. Nat. Rev. Immunol. 14, 546–558.

Hao, B., Chen, Z., Bi, B., Yu, M., Yao, S., Feng, Y., Yu, Y., Pan, L., Di, D., Luo, G., et al. (2018). Role of TLR4 as a prognostic factor for survival in various cancers: a meta-analysis. Oncotarget 9, 13088–13099.

Harris, M.A., Clark, J., Ireland, A., Lomax, J., Ashburner, M., Foulger, R., Eilbeck, K., Lewis, S., Marshall, B., Mungall, C., et al. (2004). The Gene Ontology (GO) database and informatics resource. Nucleic Acids Res. 32, D258–61.

Hayden, M.S., and Ghosh, S. (2008). Shared principles in NF-kappaB signaling. Cell 132, 344– 362.

Heilmann, S., Ratnakumar, K., Langdon, E., Kansler, E., Kim, I., Campbell, N.R., Perry, E., McMahon, A., Kaufman, C., van Rooijen, E., et al. (2015). A quantitative system for studying metastasis using transparent zebrafish. Cancer Res. 75, 4272–4282.

Hirasawa, A., Tsumaya, K., Awaji, T., Katsuma, S., Adachi, T., Yamada, M., Sugimoto, Y., Miyazaki, S., and Tsujimoto, G. (2005). Free fatty acids regulate gut incretin glucagon-like peptide-1 secretion through GPR120. Nat. Med. 11, 90–94.

Hosoda, H., Kojima, M., Matsuo, H., and Kangawa, K. (2000). Ghrelin and des-acyl ghrelin: two major forms of rat ghrelin peptide in gastrointestinal tissue. Biochem. Biophys. Res. Commun. 279, 909–913.

Hossain, F., Al-Khami, A.A., Wyczechowska, D., Hernandez, C., Zheng, L., Reiss, K., Valle, L.D., Trillo-Tinoco, J., Maj, T., Zou, W., et al. (2015). Inhibition of Fatty Acid Oxidation Modulates Immunosuppressive Functions of Myeloid-Derived Suppressor Cells and Enhances Cancer Therapies. Cancer Immunol. Res. 3, 1236–1247.

Huang, S., Rutkowsky, J.M., Snodgrass, R.G., Ono-Moore, K.D., Schneider, D.A., Newman, J.W., Adams, S.H., and Hwang, D.H. (2012). Saturated fatty acids activate TLR-mediated proinflammatory signaling pathways. J. Lipid Res. 53, 2002–2013.

Iemoto, T., Nishiumi, S., Kobayashi, T., Fujigaki, S., Hamaguchi, T., Kato, K., Shoji, H., Matsumura, Y., Honda, K., and Yoshida, M. (2019). Serum level of octanoic acid predicts the efficacy of chemotherapy for colorectal cancer. Oncol. Lett. 17, 831–842.

Ilkovitch, D., and Lopez, D.M. (2008). Immune modulation by melanoma-derived factors. Exp. Dermatol. 17, 977–985.

Itoh, Y., and Hinuma, S. (2005). GPR40, a free fatty acid receptor on pancreatic beta cells, regulates insulin secretion. Hepatol. Res. 33, 171–173.

Jao, L.-E., Wente, S.R., and Chen, W. (2013). Efficient multiplex biallelic zebrafish genome editing using a CRISPR nuclease system. Proc Natl Acad Sci USA 110, 13904–13909.

Kamei, Y., Xu, L., Heinzel, T., Torchia, J., Kurokawa, R., Gloss, B., Lin, S.C., Heyman, R.A., Rose, D.W., Glass, C.K., et al. (1996). A CBP integrator complex mediates transcriptional activation and AP-1 inhibition by nuclear receptors. Cell 85, 403–414.

Kartikasari, A.E.R., Huertas, C.S., Mitchell, A., and Plebanski, M. (2021). Tumor-Induced Inflammatory Cytokines and the Emerging Diagnostic Devices for Cancer Detection and Prognosis. Front. Oncol. 11, 692142.

Kawai, T., and Akira, S. (2007). Signaling to NF-kappaB by Toll-like receptors. Trends Mol. Med. 13, 460–469.

Kim, J.-W., and Yoon, K.-H. (2011). Glucolipotoxicity in Pancreatic β-Cells. Diabetes Metab. J. 35, 444–450.

Kim, K.H., Jo, M.S., Suh, D.S., Yoon, M.S., Shin, D.H., Lee, J.H., and Choi, K.U. (2012). Expression and significance of the TLR4/MyD88 signaling pathway in ovarian epithelial cancers. World J. Surg. Oncol. 10, 193.

Koundouros, N., and Poulogiannis, G. (2020). Reprogramming of fatty acid metabolism in cancer. Br. J. Cancer 122, 4–22.

Lancaster, G.I., Langley, K.G., Berglund, N.A., Kammoun, H.L., Reibe, S., Estevez, E., Weir, J., Mellett, N.A., Pernes, G., Conway, J.R.W., et al. (2018). Evidence that TLR4 Is Not a Receptor for Saturated Fatty Acids but Mediates Lipid-Induced Inflammation by Reprogramming Macrophage Metabolism. Cell Metab. 27, 1096–1110.e5.

Lappano, R., Sebastiani, A., Cirillo, F., Rigiracciolo, D.C., Galli, G.R., Curcio, R., Malaguarnera, R., Belfiore, A., Cappello, A.R., and Maggiolini, M. (2017). The lauric acid-activated signaling prompts apoptosis in cancer cells. Cell Death Discov. 3, 17063.

Lauterbach, M.A., Hanke, J.E., Serefidou, M., Mangan, M.S.J., Kolbe, C.-C., Hess, T., Rothe, M., Kaiser, R., Hoss, F., Gehlen, J., et al. (2019). Toll-like Receptor Signaling Rewires Macrophage Metabolism and Promotes Histone Acetylation via ATP-Citrate Lyase. Immunity 51, 997–1011.e7.

Lee, J.V., Carrer, A., Shah, S., Snyder, N.W., Wei, S., Venneti, S., Worth, A.J., Yuan, Z.-F., Lim, H.-W., Liu, S., et al. (2014). Akt-dependent metabolic reprogramming regulates tumor cell histone acetylation. Cell Metab. 20, 306–319.

Lee, J.V., Berry, C.T., Kim, K., Sen, P., Kim, T., Carrer, A., Trefely, S., Zhao, S., Fernandez, S., Barney, L.E., et al. (2018). Acetyl-CoA promotes glioblastoma cell adhesion and migration through Ca2+-NFAT signaling. Genes Dev. 32, 497–511.

Lee, J.Y., Ye, J., Gao, Z., Youn, H.S., Lee, W.H., Zhao, L., Sizemore, N., and Hwang, D.H. (2003). Reciprocal modulation of Toll-like receptor-4 signaling pathways involving MyD88 and phosphatidylinositol 3-kinase/AKT by saturated and polyunsaturated fatty acids. J. Biol. Chem. 278, 37041–37051.

Lemarié, F., Beauchamp, E., Drouin, G., Legrand, P., and Rioux, V. (2018). Dietary caprylic acid and ghrelin O-acyltransferase activity to modulate octanoylated ghrelin functions: What is new in this nutritional field? Prostaglandins Leukot. Essent. Fatty Acids 135, 121–127.

Lindsay, H., Burger, A., Biyong, B., Felker, A., Hess, C., Zaugg, J., Chiavacci, E., Anders, C., Jinek, M., Mosimann, C., et al. (2016). CrispRVariants charts the mutation spectrum of genome engineering experiments. Nat. Biotechnol. 34, 701–702.

Liu, J., Mazzone, P.J., Cata, J.P., Kurz, A., Bauer, M., Mascha, E.J., and Sessler, D.I. (2014). Serum free fatty acid biomarkers of lung cancer. Chest 146, 670–679.

Louis, P., Hold, G.L., and Flint, H.J. (2014). The gut microbiota, bacterial metabolites and colorectal cancer. Nat. Rev. Microbiol. 12, 661–672.

Maeshima, N., and Fernandez, R.C. (2013). Recognition of lipid A variants by the TLR4-MD-2 receptor complex. Front. Cell. Infect. Microbiol. 3, 3.

Matsushita, M., Fujita, K., Hayashi, T., Kayama, H., Motooka, D., Hase, H., Jingushi, K., Yamamichi, G., Yumiba, S., Tomiyama, E., et al. (2021). Gut Microbiota-Derived Short-Chain Fatty Acids Promote Prostate Cancer Growth via IGF1 Signaling. Cancer Res. 81, 4014–4026.

McDonnell, E., Crown, S.B., Fox, D.B., Kitir, B., Ilkayeva, O.R., Olsen, C.A., Grimsrud, P.A., and Hirschey, M.D. (2016). Lipids reprogram metabolism to become a major carbon source for histone acetylation. Cell Rep. 17, 1463–1472.

McGarry, J.D., and Foster, D.W. (1971). The Regulation of Ketogenesis from Octanoic Acid. Journal of Biological Chemistry 246, 1149–1159.

Meeth, K., Wang, J.X., Micevic, G., Damsky, W., and Bosenberg, M.W. (2016). The YUMM lines: a series of congenic mouse melanoma cell lines with defined genetic alterations. Pigment Cell Melanoma Res. 29, 590–597.

Menendez, J.A., and Lupu, R. (2007). Fatty acid synthase and the lipogenic phenotype in cancer pathogenesis. Nat. Rev. Cancer 7, 763–777.

Milanski, M., Degasperi, G., Coope, A., Morari, J., Denis, R., Cintra, D.E., Tsukumo, D.M.L., Anhe, G., Amaral, M.E., Takahashi, H.K., et al. (2009). Saturated fatty acids produce an inflammatory response predominantly through the activation of TLR4 signaling in hypothalamus: implications for the pathogenesis of obesity. J. Neurosci. 29, 359–370.

Mittal, D., Saccheri, F., Vénéreau, E., Pusterla, T., Bianchi, M.E., and Rescigno, M. (2010). TLR4-mediated skin carcinogenesis is dependent on immune and radioresistant cells. EMBO J. 29, 2242–2252.

Narayanan, A., Baskaran, S.A., Amalaradjou, M.A.R., and Venkitanarayanan, K. (2015). Anticarcinogenic properties of medium chain fatty acids on human colorectal, skin and breast cancer cells in vitro. Int. J. Mol. Sci. 16, 5014–5027.

Nicholas, D.A., Zhang, K., Hung, C., Glasgow, S., Aruni, A.W., Unternaehrer, J., Payne, K.J., Langridge, W.H.R., and De Leon, M. (2017). Palmitic acid is a toll-like receptor 4 ligand that induces human dendritic cell secretion of IL-1β. PLoS ONE 12, e0176793.

Oh, D.Y., Talukdar, S., Bae, E.J., Imamura, T., Morinaga, H., Fan, W., Li, P., Lu, W.J., Watkins, S.M., and Olefsky, J.M. (2010). GPR120 is an omega-3 fatty acid receptor mediating potent anti-inflammatory and insulin-sensitizing effects. Cell 142, 687–698.

Otto, C., Kaemmerer, U., Illert, B., Muehling, B., Pfetzer, N., Wittig, R., Voelker, H.U., Thiede, A., and Coy, J.F. (2008). Growth of human gastric cancer cells in nude mice is delayed by a ketogenic diet supplemented with omega-3 fatty acids and medium-chain triglycerides. BMC Cancer 8, 122.

Park, B.S., Song, D.H., Kim, H.M., Choi, B.-S., Lee, H., and Lee, J.-O. (2009). The structural basis of lipopolysaccharide recognition by the TLR4-MD-2 complex. Nature 458, 1191–1195.

Park, J.H., Vithayathil, S., Kumar, S., Sung, P.-L., Dobrolecki, L.E., Putluri, V., Bhat, V.B., Bhowmik, S.K., Gupta, V., Arora, K., et al. (2016). Fatty Acid Oxidation-Driven Src Links Mitochondrial Energy Reprogramming and Oncogenic Properties in Triple-Negative Breast Cancer. Cell Rep. 14, 2154–2165.

Pascual, G., Avgustinova, A., Mejetta, S., Martín, M., Castellanos, A., Attolini, C.S.-O., Berenguer, A., Prats, N., Toll, A., Hueto, J.A., et al. (2017). Targeting metastasis-initiating cells through the fatty acid receptor CD36. Nature 541, 41–45.

Pascual, G., Domínguez, D., Elosúa-Bayes, M., Beckedorff, F., Laudanna, C., Bigas, C., Douillet, D., Greco, C., Symeonidi, A., Hernández, I., et al. (2021). Dietary palmitic acid promotes a prometastatic memory via Schwann cells. Nature 599, 485–490.

Pavón, M.A., Arroyo-Solera, I., Céspedes, M.V., Casanova, I., León, X., and Mangues, R. (2016). uPA/uPAR and SERPINE1 in head and neck cancer: role in tumor resistance, metastasis, prognosis and therapy. Oncotarget 7, 57351–57366.

Pietrocola, F., Galluzzi, L., Bravo-San Pedro, J.M., Madeo, F., and Kroemer, G. (2015). Acetyl coenzyme A: a central metabolite and second messenger. Cell Metab. 21, 805–821.

Poitout, V., Amyot, J., Semache, M., Zarrouki, B., Hagman, D., and Fontés, G. (2010). Glucolipotoxicity of the pancreatic beta cell. Biochim. Biophys. Acta 1801, 289–298.

Poltorak, A., He, X., Smirnova, I., Liu, M.Y., Van Huffel, C., Du, X., Birdwell, D., Alejos, E., Silva, M., Galanos, C., et al. (1998). Defective LPS signaling in C3H/HeJ and C57BL/10ScCr mice: mutations in Tlr4 gene. Science 282, 2085–2088.

Qu, Q., Zeng, F., Liu, X., Wang, Q.J., and Deng, F. (2016). Fatty acid oxidation and carnitine palmitoyltransferase I: emerging therapeutic targets in cancer. Cell Death Dis. 7, e2226.

Rajput, S., Volk-Draper, L.D., and Ran, S. (2013). TLR4 is a novel determinant of the response to paclitaxel in breast cancer. Mol. Cancer Ther. 12, 1676–1687.

Rakoff-Nahoum, S., and Medzhitov, R. (2007). Regulation of spontaneous intestinal tumorigenesis through the adaptor protein MyD88. Science 317, 124–127.

Rector, R.S., Payne, R.M., and Ibdah, J.A. (2008). Mitochondrial trifunctional protein defects: clinical implications and therapeutic approaches. Adv. Drug Deliv. Rev. 60, 1488–1496.

Rocha, D.M., Caldas, A.P., Oliveira, L.L., Bressan, J., and Hermsdorff, H.H. (2016). Saturated fatty acids trigger TLR4-mediated inflammatory response. Atherosclerosis 244, 211–215.

Rogero, M.M., and Calder, P.C. (2018). Obesity, Inflammation, Toll-Like Receptor 4 and Fatty Acids. Nutrients 10.

Röhrig, F., and Schulze, A. (2016). The multifaceted roles of fatty acid synthesis in cancer. Nat. Rev. Cancer 16, 732–749.

Sakamoto, H., Koma, Y.-I., Higashino, N., Kodama, T., Tanigawa, K., Shimizu, M., Fujikawa, M., Nishio, M., Shigeoka, M., Kakeji, Y., et al. (2021). PAI-1 derived from cancer-associated fibroblasts in esophageal squamous cell carcinoma promotes the invasion of cancer cells and the migration of macrophages. Lab. Invest. 101, 353–368.

Salcedo, R., Worschech, A., Cardone, M., Jones, Y., Gyulai, Z., Dai, R.-M., Wang, E., Ma, W., Haines, D., O’hUigin, C., et al. (2010). MyD88-mediated signaling prevents development of adenocarcinomas of the colon: role of interleukin 18. J. Exp. Med. 207, 1625–1636.

Schlaepfer, I.R., Rider, L., Rodrigues, L.U., Gijón, M.A., Pac, C.T., Romero, L., Cimic, A., Sirintrapun, S.J., Glodé, L.M., Eckel, R.H., et al. (2014). Lipid catabolism via CPT1 as a therapeutic target for prostate cancer. Mol. Cancer Ther. 13, 2361–2371.

Seker, F., Cingoz, A., Sur-Erdem, İ., Erguder, N., Erkent, A., Uyulur, F., Esai Selvan, M., Gümüş, Z.H., Gönen, M., Bayraktar, H., et al. (2019). Identification of SERPINE1 as a Regulator of Glioblastoma Cell Dispersal with Transcriptome Profiling. Cancers (Basel) 11.

Shi, H., Kokoeva, M.V., Inouye, K., Tzameli, I., Yin, H., and Flier, J.S. (2006). TLR4 links innate immunity and fatty acid-induced insulin resistance. J. Clin. Invest. 116, 3015–3025.

Shi, J., Fu, H., Jia, Z., He, K., Fu, L., and Wang, W. (2016). High expression of CPT1A predicts adverse outcomes: A potential therapeutic target for acute myeloid leukemia. EBioMedicine 14, 55–64.

Sidoli, S., Bhanu, N.V., Karch, K.R., Wang, X., and Garcia, B.A. (2016). Complete Workflow for Analysis of Histone Post-translational Modifications Using Bottom-up Mass Spectrometry: From Histone Extraction to Data Analysis. J. Vis. Exp.

Sivanand, S., Rhoades, S., Jiang, Q., Lee, J.V., Benci, J., Zhang, J., Yuan, S., Viney, I., Zhao, S., Carrer, A., et al. (2017). Nuclear Acetyl-CoA Production by ACLY Promotes Homologous Recombination. Mol. Cell 67, 252–265.e6.

Sung, G.-J., Choi, H.-K., Kwak, S., Song, J.-H., Ko, H., Yoon, H.-G., Kang, H.-B., and Choi, K.-C. (2016). Targeting CPT1A enhances metabolic therapy in human melanoma cells with the BRAF V600E mutation. Int. J. Biochem. Cell Biol. 81, 76–81.

Takazawa, Y., Kiniwa, Y., Ogawa, E., Uchiyama, A., Ashida, A., Uhara, H., Goto, Y., and Okuyama, R. (2014). Toll-like receptor 4 signaling promotes the migration of human melanoma cells. Tohoku J. Exp. Med. 234, 57–65.

Thomas, S.P., and Denu, J.M. (2021). Short-chain fatty acids activate acetyltransferase p300. ELife 10.

Uhlén, M., Karlsson, M.J., Hober, A., Svensson, A.-S., Scheffel, J., Kotol, D., Zhong, W., Tebani, A., Strandberg, L., Edfors, F., et al. (2019). The human secretome. Sci. Signal. 12.

Valiente, M., Obenauf, A.C., Jin, X., Chen, Q., Zhang, X.H.-F., Lee, D.J., Chaft, J.E., Kris, M.G., Huse, J.T., Brogi, E., et al. (2014). Serpins promote cancer cell survival and vascular co-option in brain metastasis. Cell 156, 1002–1016.

Vanden Berghe, W., De Bosscher, K., Boone, E., Plaisance, S., and Haegeman, G. (1999). The nuclear factor-kappaB engages CBP/p300 and histone acetyltransferase activity for transcriptional activation of the interleukin-6 gene promoter. J. Biol. Chem. 274, 32091–32098.

Veglia, F., Tyurin, V.A., Blasi, M., De Leo, A., Kossenkov, A.V., Donthireddy, L., To, T.K.J., Schug, Z., Basu, S., Wang, F., et al. (2019). Fatty acid transport protein 2 reprograms neutrophils in cancer. Nature 569, 73–78.

Vinci, M., Gowan, S., Boxall, F., Patterson, L., Zimmermann, M., Court, W., Lomas, C., Mendiola, M., Hardisson, D., and Eccles, S.A. (2012). Advances in establishment and analysis of three-dimensional tumor spheroid-based functional assays for target validation and drug evaluation. BMC Biol. 10, 29.

Vinci, M., Box, C., and Eccles, S.A. (2015). Three-dimensional (3D) tumor spheroid invasion assay. J. Vis. Exp. e52686.

Wang, C., Wang, J., Chen, K., Pang, H., Li, X., Zhu, J., Ma, Y., Qiu, T., Li, W., Xie, J., et al. (2020). Caprylic acid (C8:0) promotes bone metastasis of prostate cancer by dysregulated adipo-osteogenic balance in bone marrow. Cancer Sci. 111, 3600–3612.

Wang, K., Wang, J., Wei, F., Zhao, N., Yang, F., and Ren, X. (2017). Expression of TLR4 in Non-Small Cell Lung Cancer Is Associated with PD-L1 and Poor Prognosis in Patients Receiving Pulmonectomy. Front. Immunol. 8, 456.

Wang, T., Fahrmann, J.F., Lee, H., Li, Y.-J., Tripathi, S.C., Yue, C., Zhang, C., Lifshitz, V., Song, J., Yuan, Y., et al. (2018). JAK/STAT3-Regulated Fatty Acid β-Oxidation Is Critical for Breast Cancer Stem Cell Self-Renewal and Chemoresistance. Cell Metab. 27, 136–150.e5.

Wellen, K.E., Hatzivassiliou, G., Sachdeva, U.M., Bui, T.V., Cross, J.R., and Thompson, C.B. (2009). ATP-citrate lyase links cellular metabolism to histone acetylation. Science 324, 1076– 1080.

White, R.M., Sessa, A., Burke, C., Bowman, T., LeBlanc, J., Ceol, C., Bourque, C., Dovey, M., Goessling, W., Burns, C.E., et al. (2008). Transparent adult zebrafish as a tool for in vivo transplantation analysis. Cell Stem Cell 2, 183–189.

Wong, S.W., Kwon, M.-J., Choi, A.M.K., Kim, H.-P., Nakahira, K., and Hwang, D.H. (2009). Fatty acids modulate Toll-like receptor 4 activation through regulation of receptor dimerization and recruitment into lipid rafts in a reactive oxygen species-dependent manner. J. Biol. Chem. 284, 27384–27392.

Xhemalce, B., and Kouzarides, T. (2010). A chromodomain switch mediated by histone H3 Lys 4 acetylation regulates heterochromatin assembly. Genes Dev. 24, 647–652.

Yang, J., Brown, M.S., Liang, G., Grishin, N.V., and Goldstein, J.L. (2008). Identification of the acyltransferase that octanoylates ghrelin, an appetite-stimulating peptide hormone. Cell 132, 387–396.

Yusuf, N., Nasti, T.H., Long, J.A., Naseemuddin, M., Lucas, A.P., Xu, H., and Elmets, C.A. (2008). Protective role of Toll-like receptor 4 during the initiation stage of cutaneous chemical carcinogenesis. Cancer Res. 68, 615–622.

Zhang, L., Han, L., He, J., Lv, J., Pan, R., and Lv, T. (2020). A high serum-free fatty acid level is associated with cancer. J. Cancer Res. Clin. Oncol. 146, 705–710.

Zhang, M., Di Martino, J.S., Bowman, R.L., Campbell, N.R., Baksh, S.C., Simon-Vermot, T., Kim, I.S., Haldeman, P., Mondal, C., Yong-Gonzales, V., et al. (2018a). Adipocyte-Derived Lipids Mediate Melanoma Progression via FATP Proteins. Cancer Discov. 8, 1006–1025.

Zhang, Y., Wang, Y., Wang, X., Ji, Y., Cheng, S., Wang, M., Zhang, C., Yu, X., Zhao, R., Zhang, W., et al. (2018b). Acetyl-coenzyme A acyltransferase 2 promote the differentiation of sheep precursor adipocytes into adipocytes. J. Cell. Biochem.

